# Modelling the transmission and spread of yellow fever in forest landscapes with different spatial configurations

**DOI:** 10.1101/2023.11.11.566684

**Authors:** Antônio Ralph Medeiros-Sousa, Martin Lange, Luis Filipe Mucci, Mauro Toledo Marrelli, Volker Grimm

## Abstract

Yellow fever (YF) is a major public health issue in tropical and subtropical areas of Africa and South America. The disease is caused by the yellow fever virus (YFV), an RNA virus transmitted to humans and other animals through the bite of infected mosquitoes (Diptera: Culicidae). In Brazil and other South American countries, YFV is restricted to the sylvatic cycle, with periodic epizootic outbreaks affecting non-human primate (NHP) populations and preceding the emergence of human infections in areas close to forests. In recent epizootic-epidemic waves, the virus has expanded its range and spread across highly fragmented landscapes of the Brazilian Atlantic coast. Empirical evidence has suggested a possible relationship between highly fragmented areas, increased risk of disease in NHP and humans, and easier permeability of YFV through the landscape. Here, we present a hybrid compartmental and network-based model to simulate the transmission and spread of YFV in forest landscapes with different spatial configurations (forest cover and edge densities) and apply the model to test the hypothesis of faster virus percolation in highly fragmented landscapes. The model was parameterized and tested using the pattern-oriented modelling approach. Two different scenarios were simulated to test variations in model outputs, a first where the landscape has no influence on model parameters (default) and a second based on the hypothesis that edge density influences mosquito and dead-end host abundance and dispersal (landscape-dependent). The model was able to reproduce empirical patterns such as the percolation speed of the virus, which presented averages close to 1 km/day, and provided insights into the short persistence time of the virus in the landscape, which was approximately three months on average. When assessing the speed of virus percolation across landscapes, it was found that in the default scenario virus percolation tended to be faster in landscapes with greater forest cover and lower edge density, which contradicts empirical observations. Conversely, in the landscape-dependent scenario, virus percolation was faster in landscapes with high edge density and intermediate forest cover, supporting empirical observations that highly fragmented landscapes favour YFV spread. The proposed model can contribute to the understanding of the dynamics of YFV spread in forested areas, with the potential to be used as an additional tool to support prevention and control measures. The potential applications of the model for YFV and other mosquito-borne diseases are discussed.

## 1. Introduction

Yellow fever (YF) is an infectious disease whose etiological agent is the YFV (yellow fever virus), an RNA virus belonging to the Flaviviridae family (Zanotto et al., 1996). The virus is transmitted to humans and other hosts by the bites of infected mosquitoes (Diptera: Culicidae). In humans, the clinical manifestations range from mild fever to severe liver disease with bleeding and yellowing skin (jaundice). Among patients who develop the severe form of yellow fever, there is a high lethality rate, ranging from 20 to 60% (Monath and Vasconcelos, 2015). Despite the availability of a safe and effective vaccine since the 1930s, yellow fever still poses a significant public health concern in tropical and subtropical regions of Africa and South America. The global burden was estimated at 109,000 severe cases and 51,000 deaths in 2018 (WHO, 2018; Gaythorpe et al., 2021).

Urban epidemics of yellow fever frequently occurred in South America until the mid-1940s, having been eliminated from cities after mass vaccination campaigns and combating the urban vector, the mosquito *Aedes aegypti* (Franco, 1969). Since then, the circulation of the virus on the continent has been restricted to forest areas, the so-called sylvatic yellow fever (SYF), with mosquitoes of the *Haemagogus* and *Sabethes* genera as the main vectors (Hervé et al., 1986; Abreu et al., 2019a). Every year cases and deaths of humans and non-human primates (NHP) have been reported in the Amazon region, which is considered an endemic and enzootic area for the virus (Bryant et al., 2003; Barret and Higgs, 2007).

Dispersion events of YFV from endemic to non-endemic extra-Amazonian regions of Brazil occur at least once every decade, causing epizootic and epidemic waves towards the East and South territories of the country (Vasconcelos et al. 2001; Monath & Vasconcelos, 2015). The SYF re-emergence in South America between 2014 – 2020 expanded the transmission to areas with no vaccine recommendation in the Atlantic Coast of southeastern Brazil, spreading through highly fragmented landscapes and reaching forests in the vicinity of densely populated areas, as the cities of São Paulo and Rio de Janeiro, significantly increasing the risk of re-urbanization of the disease (Possas et al., 2018; Brazilian Ministry of Health, 2019). A total of 2,216 NHP deaths and 779 human deaths from 2,273 human cases were reported in Brazil from 2014 to June 2020, of which 98% occurred in the southeastern states (Brazilian Ministry of Health, 2021).

The spread of YFV in the Atlantic Forest was facilitated through ecological corridors created by the remaining forest fragments (Brazilian Ministry of Health, 2019). These areas serve as structural pathways and habitats for various new-world NHP, including howler monkeys, marmosets, and capuchin monkeys, as well as vector mosquitoes such as *Haemagogus* spp. and *Sabethes* spp. (Possas et al., 2018). Recent studies suggest that highly fragmented forested areas are more prone to YFV (Prist et al., 2022, Wilk-da-Silva et al., 2022), increasing the risk of SYF disease occurrence in monkeys and humans (Ilacqua et al., 2021). A possible mechanism for this is a reduction in the biodiversity of non-competent hosts and increased mosquito dispersal between forest fragments separated by open areas. These studies were based on empirical data from epizootic waves that occurred in Southeast Brazil between 2017 and 2019.

Such observations have led to the hypothesis that high edge density and good connectivity between fragments, characteristics commonly observed in landscapes with intermediate levels of forest cover (∼30 – 70%) (Ilacqua et al., 2021), facilitate the spread of the virus. The arguments are that increased edge areas found in these environments present: 1) lower relative humidity and higher temperature compared to the interior of forests, which favours a greater abundance of potential YFV vectors, in particular *Hg. leucocelaenus* that shows good adaptation to degraded forest environments (Camargo-Neves et al., 2005; Magnano et al., 2015; Hendy et al., 2020); 2) fewer suitable sites for oviposition, increasing the need for long-distance flights for mosquitoes, in addition to greater exposure to strong winds, which can increase mosquito dispersal capacity (Magnano et al., 2015; Almeida et al. al., 2019); 3) lower diversity of non-competent vertebrate hosts to transmit viruses (dead-end hosts), reducing the dilution effect of the pathogen in the environment (see Kessing et al., 2010).

Understanding the impact of landscape fragmentation on pathogen spread is crucial for assessing the risk to human health (Allan et al., 2003, Gottwalt, 2013). In the case of SYF, it would also allow the identification of priority areas for prevention and the allocation of resources for vaccination campaigns (Ilacqua et al., 2021, Prist et al., 2022). However, obtaining empirical data necessary for a better understanding of the SYF dynamics is difficult due to the numerous variables and species involved in the transmission cycle. This difficulty leads to high costs and time consumption, making it even impossible to gain comprehensive insights into the diverse aspects of the disease. For these reasons, modelling the transmission and dispersion dynamics of the virus between host and vector is an alternative to test and propose new hypotheses on the ecology and epidemiology of the disease, with the ability to identify priority issues to be investigated in empirical studies and to predict risk situations. Although a variety of modelling approaches have been used to study the ecological and epidemiological aspects of SYF (some examples are Moreno et al., 2015, Ribeiro et al., 2015, Hamlet et al., 2021, Cunha et al., 2022, Li et al., 2022, Wilk-da-Silva et al., 2022), to date there are no models that dynamically simulate how vector, host and landscape characteristics, and the interaction between these elements, determine the transmission and spread of YFV in forest areas.

In this study, we present a hybrid compartment- and network-based dynamical model to simulate the transmission and spread of the SYF in fragmented landscapes. The objective is 1) to describe in detail the structure of the model, 2) to evaluate the capacity of the model to reproduce empirically observed patterns and 3) to demonstrate a first theoretical application of the model, evaluating in which transmission scenarios there is support for the hypothesis that highly fragmented landscapes favour the permeability of the virus. The potential application of the model to investigate different issues involving SYF and other mosquito-borne diseases is discussed.

## 2. Methods

### 2.1. Model description

This model description follows the ODD (Overview, Design concepts, Details) protocol for describing agent-based models (Grimm et al. 2006, 2010, 2020). Here we present a summarized description of the model. A full and detailed ODD protocol is presented in Supplementary Information S1. The model was implemented in NetLogo (Wilensky 1999), version 6.3.0 and the current version is available at https://github.com/aralphms/YFVLandModel.

The overall *purpose* of the model is to represent the transmission dynamics and propagation of the yellow fever virus (YFV) across fragmented landscapes. The within-fragment dynamics is based on the local agent-based model for YFV transmission developed by Medeiros-Sousa et al. (2022), which for run-time reasons was aggregated into a compartment model. Specifically, we are addressing the following questions: Do highly fragmented landscapes facilitate the spread of YFV during epizootic waves? The main hypothesis is that edge areas of forests have higher abundances of the main vector, mosquitoes, and less diversity of dead-end hosts, which increases the contact rates between vectors and monkeys and facilitates virus amplification and propagation in the landscape. In addition, fragmented areas may facilitate the dispersal of mosquitoes for long distances looking for blood sources and breeding sites or being carried by the wind. Thus, high permeability to the virus is expected in highly fragmented landscapes with an intermediate degree of forest cover since these areas tend to present high edge density, support large populations of monkeys and mosquitoes, have a lower diversity of dead-end hosts, and have good connectivity between fragments allowing higher dispersal of mosquitoes and hosts.

To consider our model realistic enough for its purpose, we use *patterns* in YFV spread speed (average, variation, and seasonal differences), the YFV persistence in the landscape, the mortality of main hosts (howler-monkeys), and the observed proportion of infected mosquitoes during the peak of epizootic events.

The model includes the following *entities*: patches, forest fragments, landscape units, nodes, and links (Figure 1).

**Figure 1.**
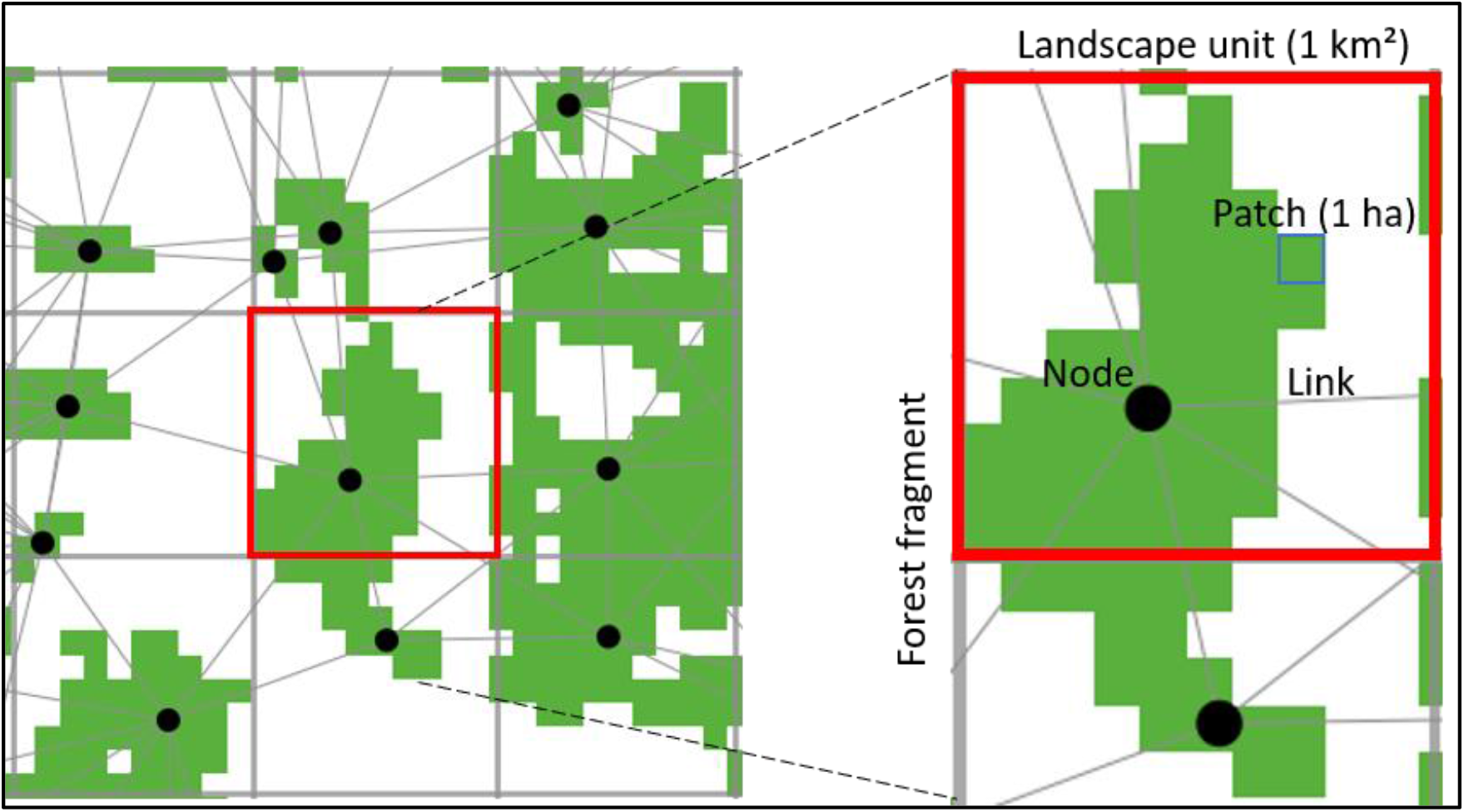
Example landscape to illustrate the model entities. The zoomed forest fragment has part of its patches located in one landscape unit (red square) and another part located in a second landscape unit. Thus, the model assigns two nodes (black circles) for the fragment. Each node has links to its neighbours.

Patches are the discrete spatial units of 100x100 m² (one hectare) defining the modelled landscape and can be of two types: forest or non-forest. Forest fragments are formed by a set of patches of the type ‘forest’. Landscape units are defined by a grid of larger spatial units of 1 km² each. A landscape unit can contain none, one or multiple forest fragments. Large forest fragments can be located in two or more landscape units. Landscape units are used to make modelling movement between fragments more efficient by assigning, for forest fragments that are larger than a landscape unit, all mosquitos within different units to different “nodes” (see below).

Nodes are abstract immobile entities that represent, for a forest fragment, the different reproductive and epidemic states of both mosquitoes and vertebrate hosts. Nodes were introduced to avoid a spatially explicit representation of vector and host dynamics within forest fragments, which would have led to too high computation times. Each node contains four different lists that represent mosquitoes (one list for adults and two for immatures) and host populations (one list that includes monkeys, alternative hosts, and dead-end hosts). The list items (hereafter compartments) indicate the number of individuals in the different reproductive and epidemic states in the populations. Links indicate which nodes are spatially linked in the grid and determine the exchange of vectors and hosts between nodes. The state variables characterizing these entities are listed in Table 1.

**Table 1.**
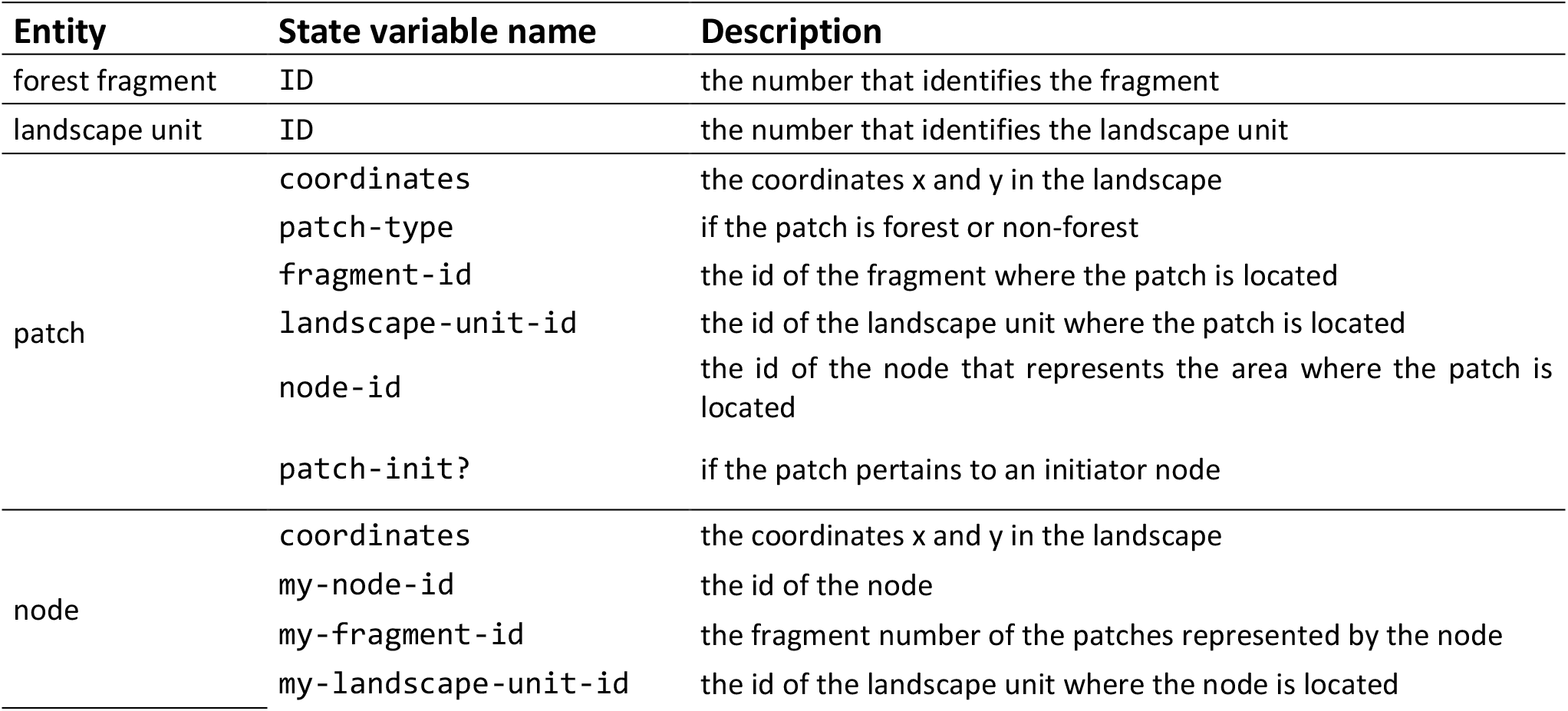
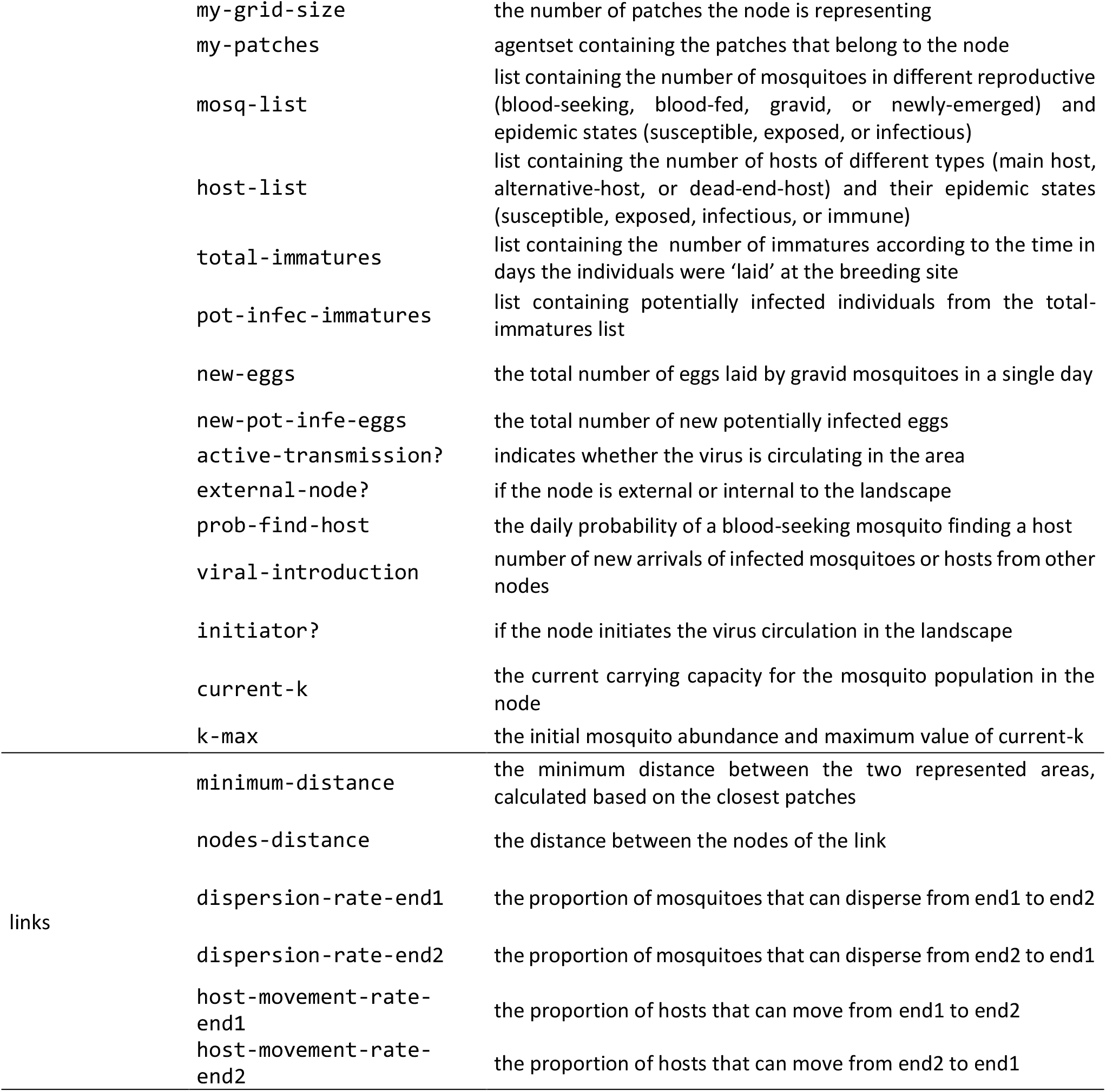
Description of the state variables assigned to each entity of the model. The structure and rationale of the compartmental structure involving the four lists characterizing a node are presented below.

As for *spatial* and *temporal* resolution and extent: The landscape consists of 121 x 121 patches, representing an area of approximately 144 km². The time step of the model represents one day, and the simulation comprises the period from the emergence of the virus to its disappearance in the landscape or when the simulation reaches 1825 days (5 years of 365 days).

The model includes a compartment-based structure that allows simulation of the transmission dynamics in fragmented landscapes with thousands or even millions of mosquitoes. In this model structure, instead of individual agents representing mosquitoes or hosts, different compartments represent the number of individuals in a given state or combination of states and for a given node. Three compartment-based structures are used in the model, one for adult mosquitoes, a second for immature forms, and a third for hosts. The compartment-based structure for mosquitoes is a matrix that represents SEI states for viral infection in the rows (susceptible, exposed, and infectious), and four reproductive states (blood-seeking, blood-fed, gravid, and newly-emerged), in the columns (figure 2A). The immature forms are represented by compartments that form the breeding site structure in the model. Each compartment indicates the number of individuals and represents the time in days since these individuals were ‘laid’ at the breeding site (figure 2B). In addition, a second list is used to count the number of potentially infected individuals (those potentially infected via transovarial transmission) in each compartment of the immature list. Host populations can be represented by SEIR states for viral infection in the rows (susceptible, exposed, infectious, and recovered/removed) and by different types of hosts in the columns. Three types of hosts are considered: main hosts, alternative hosts, and dead-end hosts (figure 2C). Matrices of *m* × *n* states can be used to represent the compartments and transitions, as illustrated in figure 2.

**Figure 2:**
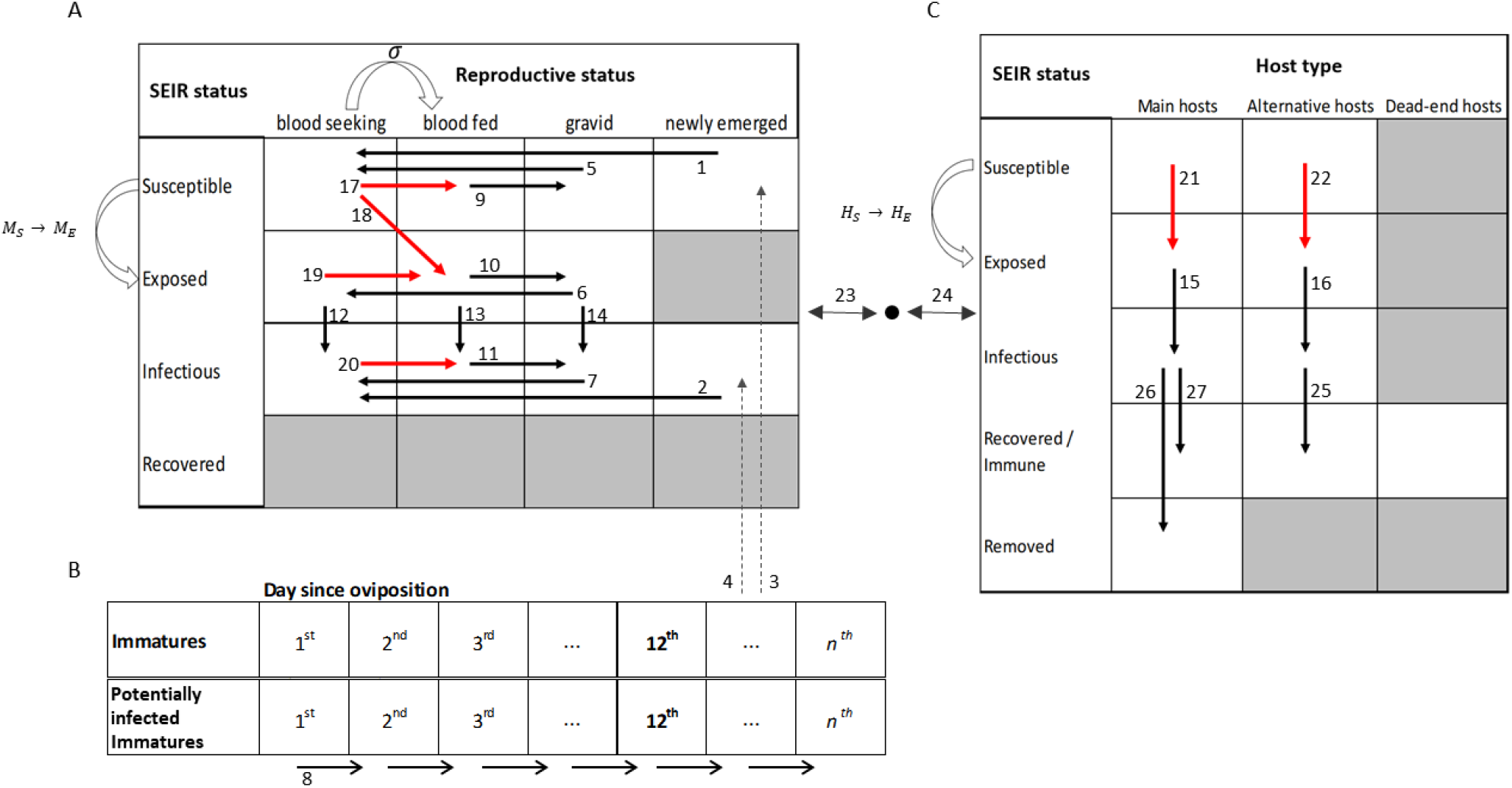
Representation of the compartment-based structure of the model. A – compartments for adult mosquitoes showing possible combinations (white compartments) between SEI and reproductive status. B – Compartments for immature forms representing the number of individuals on consecutive days after oviposition. C – Compartments for hosts showing SEIR states. The arrows represent the possible compartment transitions and the numbers represent the order in which the transitions occur in the model (the same numbers are used, e.g. “Transition 1”, as comments in the NetLogo program so that the corresponding code can easily be found). Red arrows indicate transitions that require host-vector interactions (σ) and change the state of mosquitoes from blood-seeking to blood-feeding (curved arrow on top of A). Infective interactions (curved arrows to the left sides of A and C) make susceptible individuals change to the exposed compartment (*MS* → *ME* or *HS* → *HE*). Dashed lines represent the transference of individuals from the immature to adult compartments. Double arrows represent the transference of individuals (coming from different compartments) from and to a neighbour node (black circle).

The model has procedures (*processes*) that are repeated at every *time step*. Table 2 presents a summarized description of these procedures, along with the entities executing transitions associated with each one.

**Table 2.**
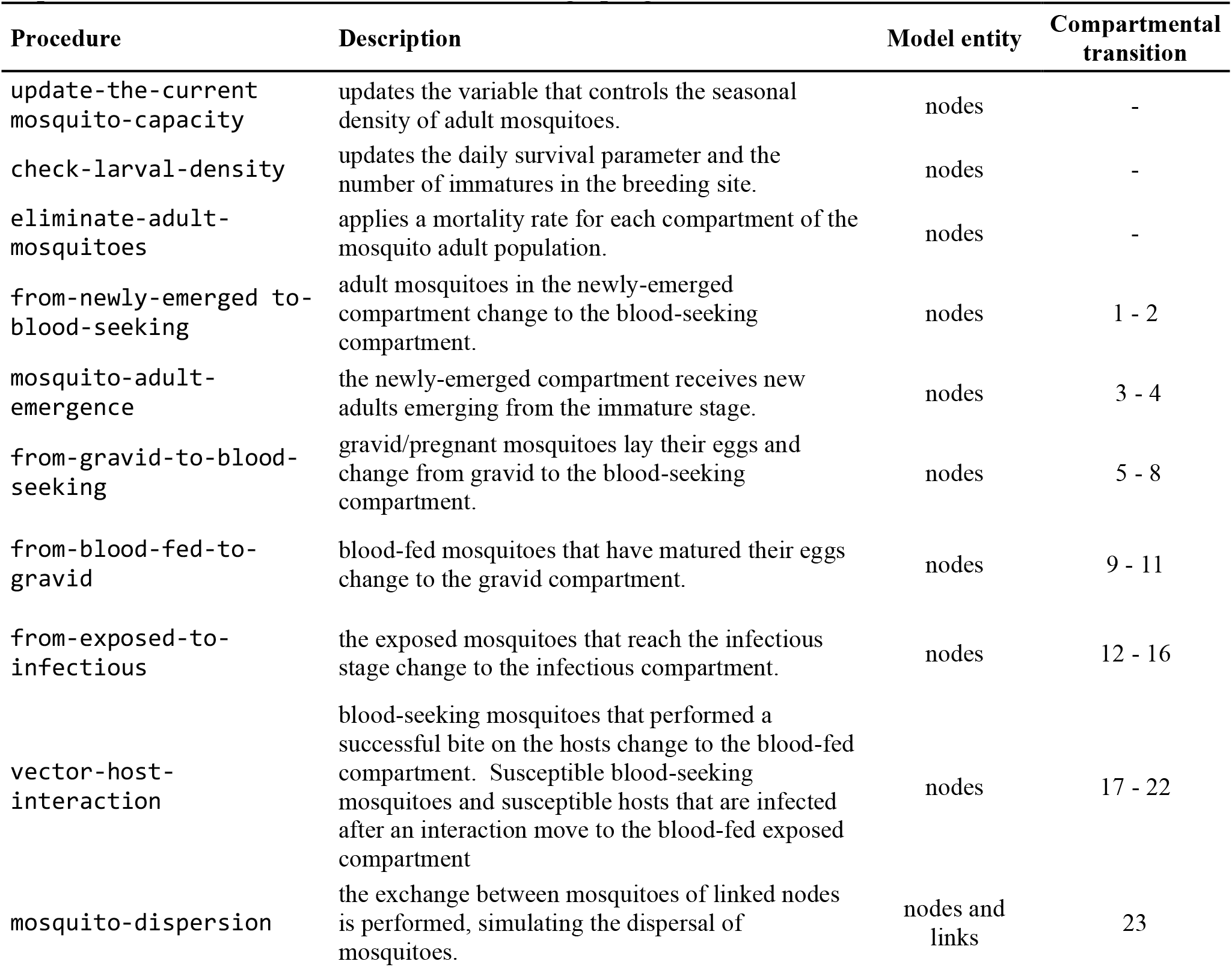
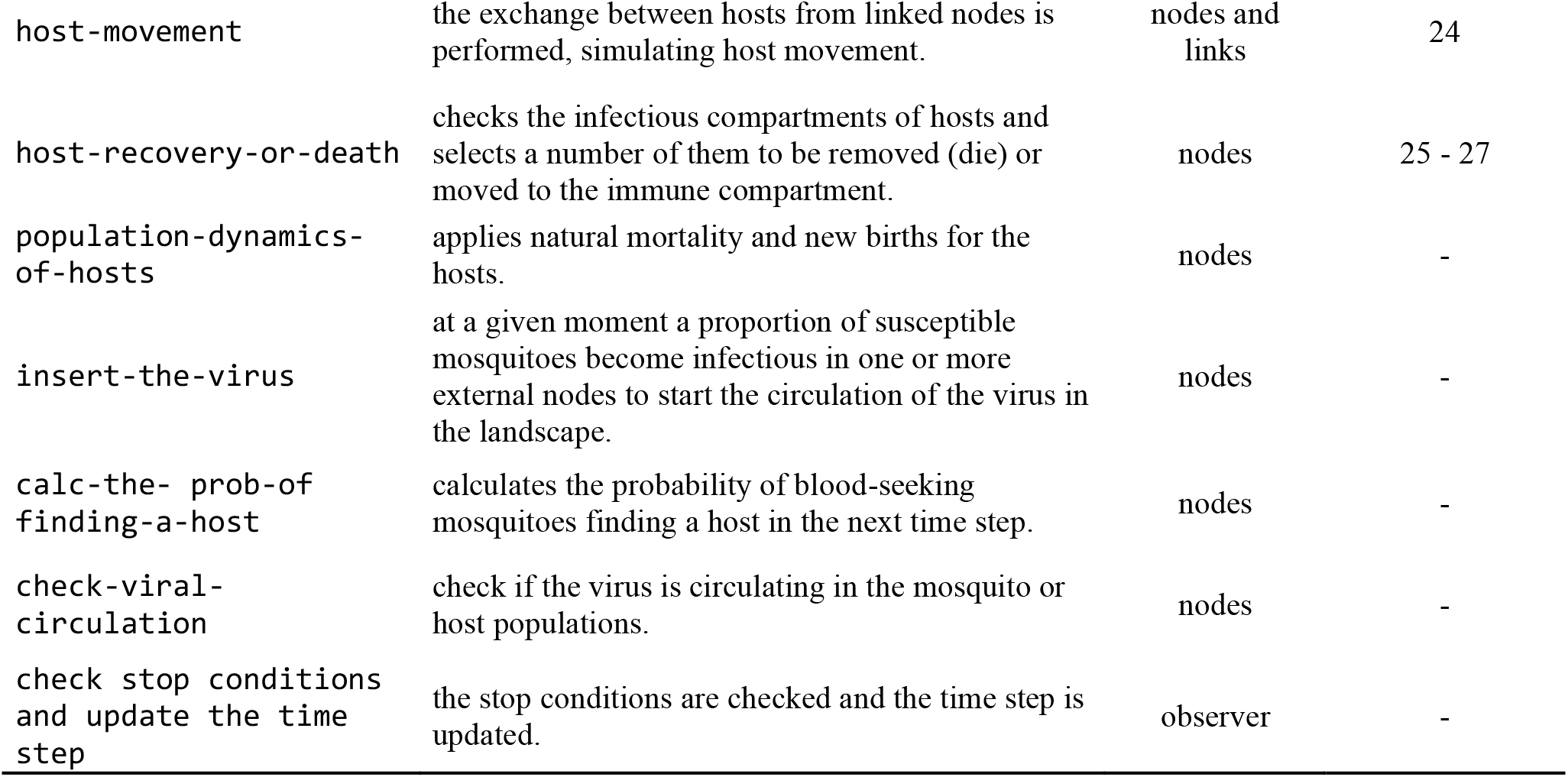
Procedures executed by the model at each time step according to the order they are called. The corresponding description, the model entity executing the procedure, and the index of the compartmental transition (as illustrated in figure 2) are shown. The same procedure names are also used in the NetLogo program.

The most important *design concept* of the model is to represent how the YFV spreads through a landscape. At the local, within fragment level, the transmission of the YFV occurs when an infectious mosquito (a female individual) feeds on the blood of a susceptible vertebrate host or, conversely, when a susceptible mosquito becomes infected by feeding on the blood of an infectious host. At the landscape level, the movement of hosts through their territories and forest corridors and the dispersal of mosquitoes over short and long distances (via active flight or carried by the wind) allows the spread of YFV to different fragments. The spread and persistence of the virus in the landscape, as well as the mortality of hosts, are patterns that emerge from the different characteristics and behaviours assigned to the model entities and can be influenced by landscape configurations.

For model *initialization,* we use vector layers of real landscapes with an extent of 12 x 12 km (see section 2.3). A second input shapefile containing a grid of 144 grid cells of 1 km² is superimposed on the landscape, dividing it into several small landscape units where the model nodes are created. The total number of nodes created in the grid is variable and depends on 1) the landscape configuration (quantity, size, and shape of fragments), and 2) a function to merge small nodes into large ones. Nodes are merged if they have less than a given number of patches and since the nodes being merged belong to the same forest fragment. All mosquitoes and hosts are initialized in the susceptible compartments. The initial day of virus introduction is assigned during initialization and is defined to simulate the YFV emergence and spread during a given season of the year.

To simulate the transmission and propagation dynamic of YFV, at a certain point in the simulation, the virus is introduced in the landscape. Two parameters control the virus insertion in the landscape: 1) the start-YFV-circulation that indicates the day the first infectious individuals will be introduced in the landscape, and 2) the YFV- initial-burden, that defines the proportion of infectious mosquitoes introduced per selected node. Five nodes spatially clustered and close to the border of the simulated ‘world’ are randomly selected and for each selected node a proportion of susceptible mosquitoes, defined by the parameter YFV-initial-burden, is moved to the compartment of infectious individuals.

Key *processes* in the model are the vector-host interaction (within fragment transmission dynamics) and the movement of mosquitoes and hosts between nodes (viral spread in the landscape). To simulate the interaction between mosquitoes and hosts in the model, we calculate four values:

1. the mosquito interaction probability with hosts (σ), calculated as the probability of a blood-seeking mosquito finding a host times the daily biting success.
2. the daily number of newly exposed/infected mosquitoes (*M*_*nE*_):

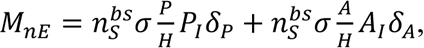

which includes the number of susceptible blood-seeking mosquitoes (*n*^*bs*^_*s*_), the proportion of main hosts and alternative hosts among the total number of hosts (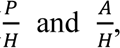 respectively), the proportion of infectious individuals among the main hosts (*P*_*I*_) and alternative hosts (*A*_*I*_), and the transmission competence of infectious main hosts (δ_*P*_) and infectious alternative hosts (δ_*A*_).
3. the daily number of newly exposed/infected main hosts (*P*_*nE*_):

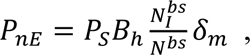

which includes, the number of susceptible main hosts (*P*^*S*^_*s*_), the infectious blood-seeking mosquitos over the total blood-seeking 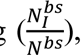, the daily number of bites per host (*B*_*h*_ = 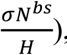, and the transmission competence of infectious mosquitoes (δ*_m_*).
4. the daily number of newly exposed/infected alternative hosts (*A*_*nE*_), calculated like *P*_*nE*_after replacing the variable *P*_*S*_ by *A*_*S*_ (susceptible alternative hosts).

After obtaining these values for mosquitoes and hosts, they are used to generate the “real” (stochastic) daily numbers of exposed individuals, using a random-Poisson distribution (λ = *M*_*nE*_; λ = *P*_*nE*_; λ = *A*_*nE*_). Finally, the values are added to their respective new compartments and subtracted from their old compartments.

We assume that the dispersal of mosquitoes is a function of the minimum distance between two linked node areas and their proportional size in relation to a landscape unit. The dispersal rate of mosquitoes (*M*_*disp*_) between two linked nodes is calculated by the link during the model initialization using the following equations:

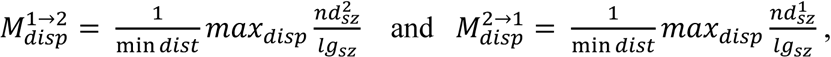

where, *M*^1→2^_*disp*_ is the dispersal rate from node 1 to node 2, *M*^2→1^_*disp*_ is the dispersal rate from node 2 to node 1, min *dist* is the minimum distance between the two forest areas represented by the two linked nodes, *max*_*disp*_ is the parameter for the maximum dispersal of mosquitoes between two nodes (max-prop-mosq-dispersal, see Table 3), *nd*^1^_*sz*_ and *nd*^2^_*sz*_ are the size in patches of the forest areas represented by the two linked nodes, and *lg*_*sz*_ is the size in patches of a landscape unit (100 patches in the current model) To determine the number of mosquitoes that will move between linked nodes, the number of individuals in each compartment of nodes 1 and 2 is multiplied by *M*^1→2^ or *M*^1→2^, respectively. After this, each node subtracts the number of dispersed individuals from their respective compartments and adds the number of migrants coming from the linked node. If an exposed or infectious individual arrives at the node, a new virus emergence event is reported.

**Table 3.**
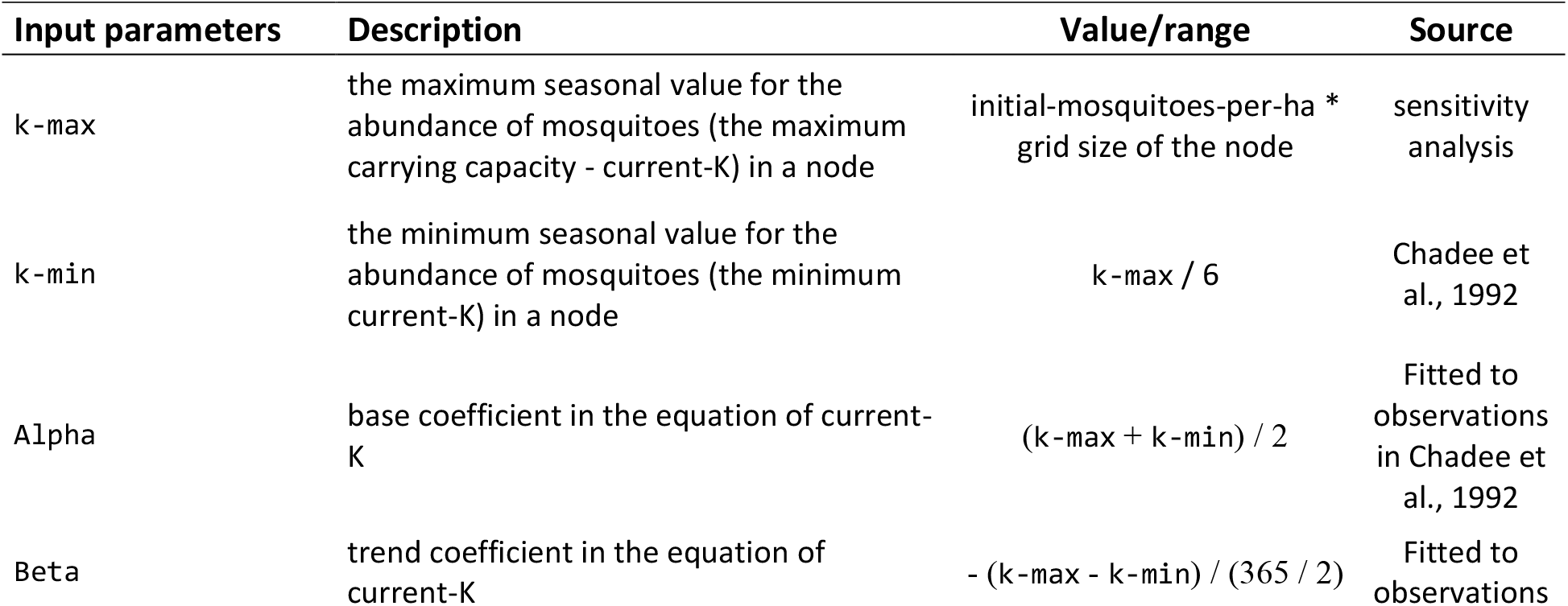
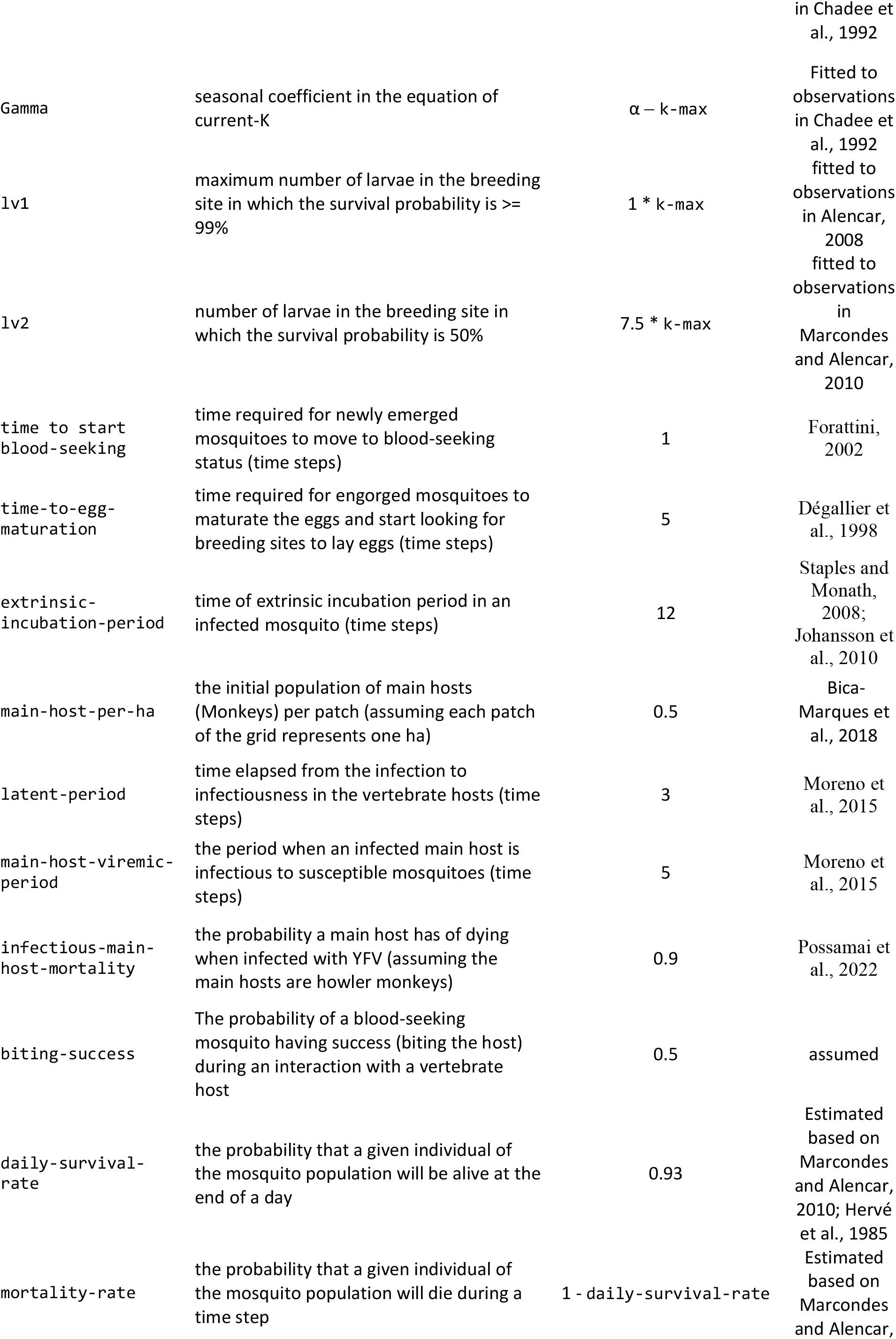
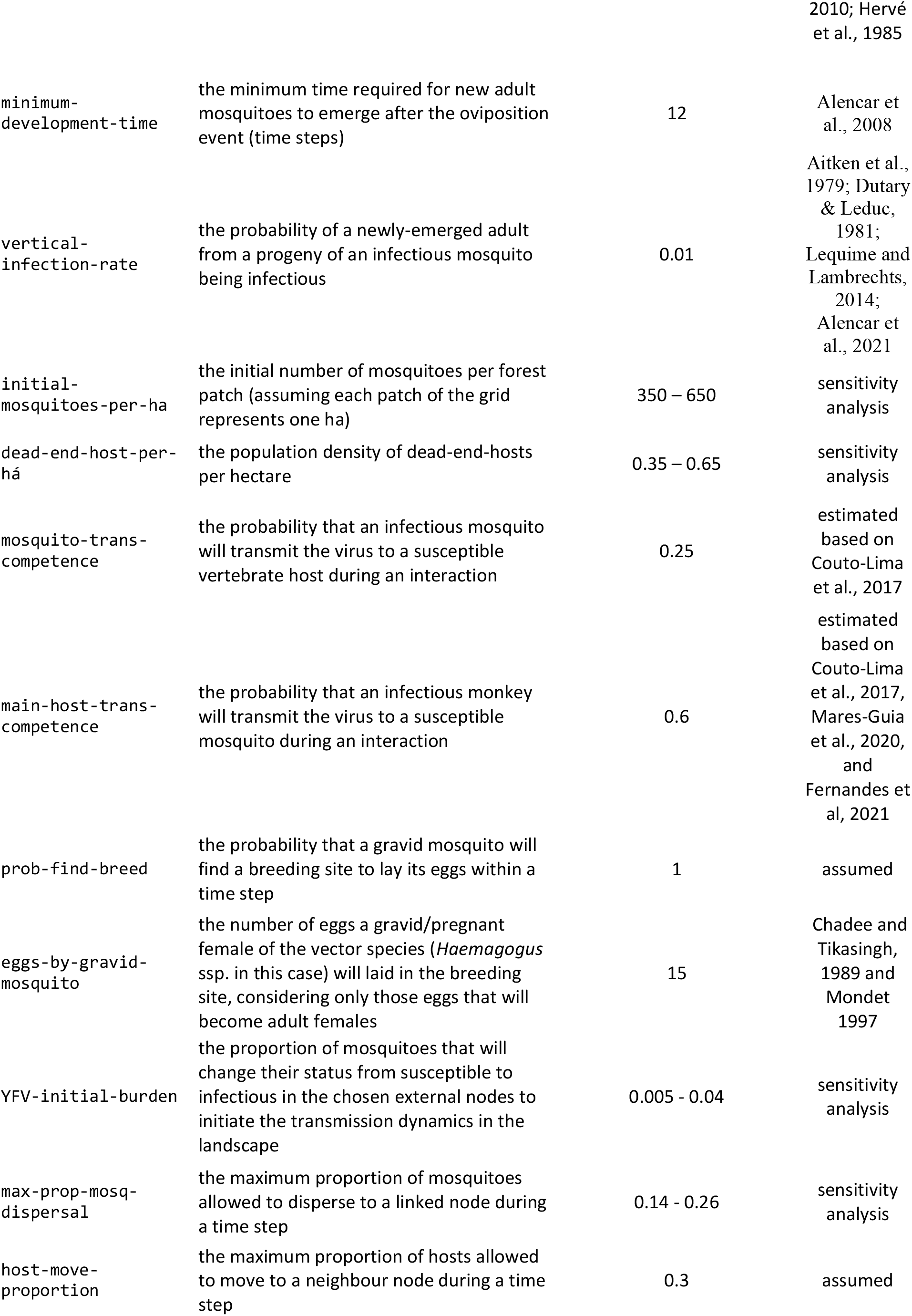
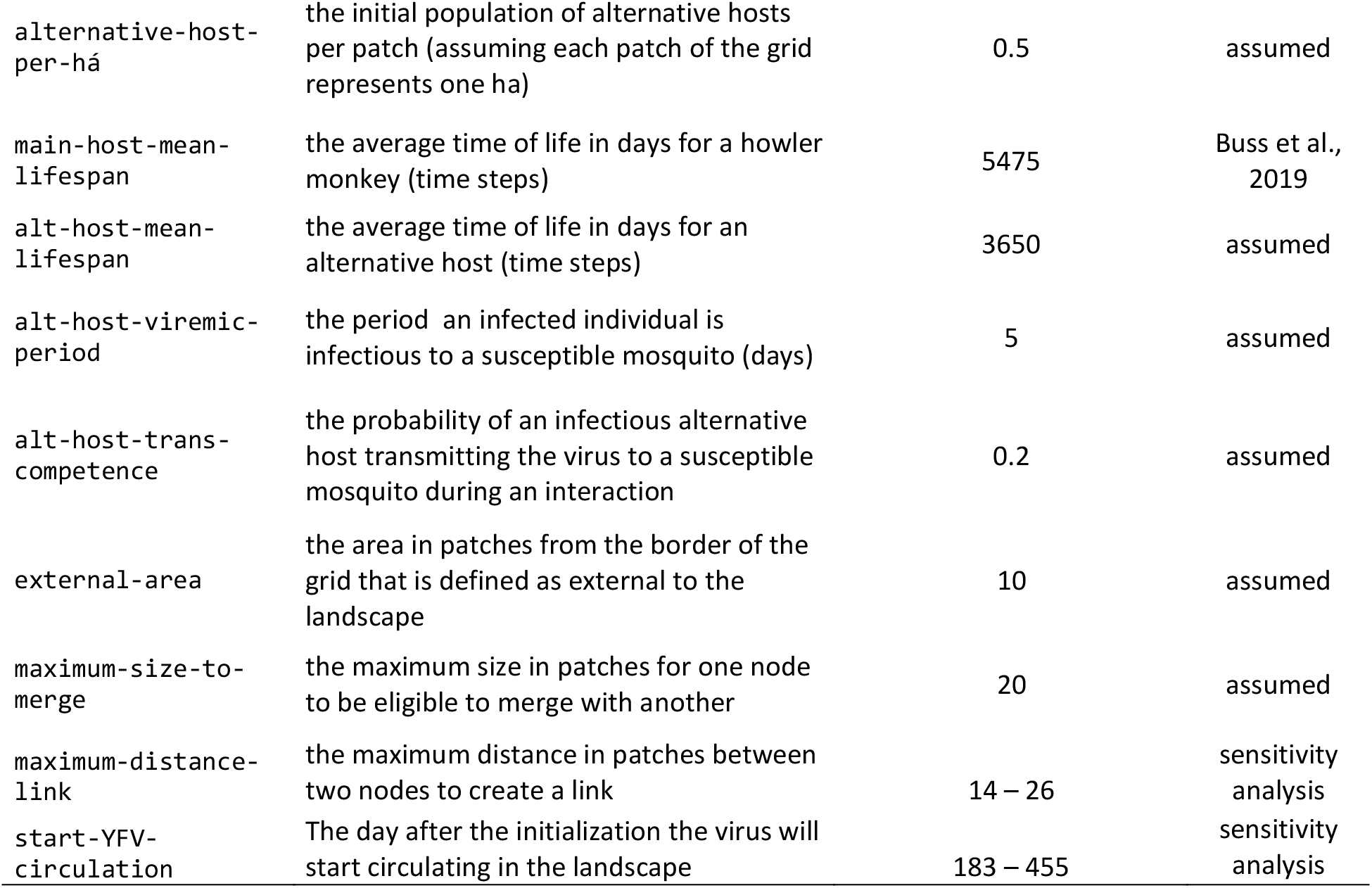
Model parameters and their values or range of values assigned during the model simulations. The description and assignment sources are also presented.

Since howler monkeys tend to not move through extensive deforested areas to migrate from one fragment to another (Hervé, 1986; Bicca-Marques and Freitas, 2010), the model assumes the movement of main hosts in the landscape is limited to linked nodes with a minimum distance equal to or lower than 2 patches (200 m). For simplicity, the same dispersion limitation was assumed for alternative and dead-end hosts. The entire procedure is performed using the same sequence described for mosquito dispersion, but replace the parameter *max*_*disp*_ by the parameter host-movement and then calculates *M*^1→2^_*disp*_ and *M*^2→1^_*disp*_ (now defined as ‘host movement rate between nodes’).

### 2.2. Model parameterization and validation

The model was parameterized and tested following the pattern-oriented modelling approach (Wiegand et al., 2004, Grimm et al., 2005, Grimm and Railsback, 2012). To assign the model’s parameter values (38 parameters, Table 3) data were obtained from observational and experimental studies, calibration, and plausible assumptions based on expert knowledge. The values of 16 parameters were assigned from field or laboratory studies, of which 12 were directly assigned values from the literature and 4 were estimated from available information. For the parameters for which there is no information in the literature, 10 were guesstimated based on discussion and consensus among experts involved in the study, five were fitted to reproduce empirical patterns related to them (see Supplementary information S1: section 7.1, text S2, and figure S1), and another seven were submitted to sensitivity analyses because they were directly related to the study hypothesis (see section 2.3).

To assess the ability of the model to reproduce empirical patterns, the final parameterization was used in replicated simulation experiments and the outputs were compared to empirical observations for YFV dynamics in fragmented landscapes. The model outputs were compared quantitatively and qualitatively to seven empirical patterns describing the virus movement in the landscape (4 patterns), persistence (1 pattern), and local within fragment patterns (2 patterns) (table 4).

**Table 4.**
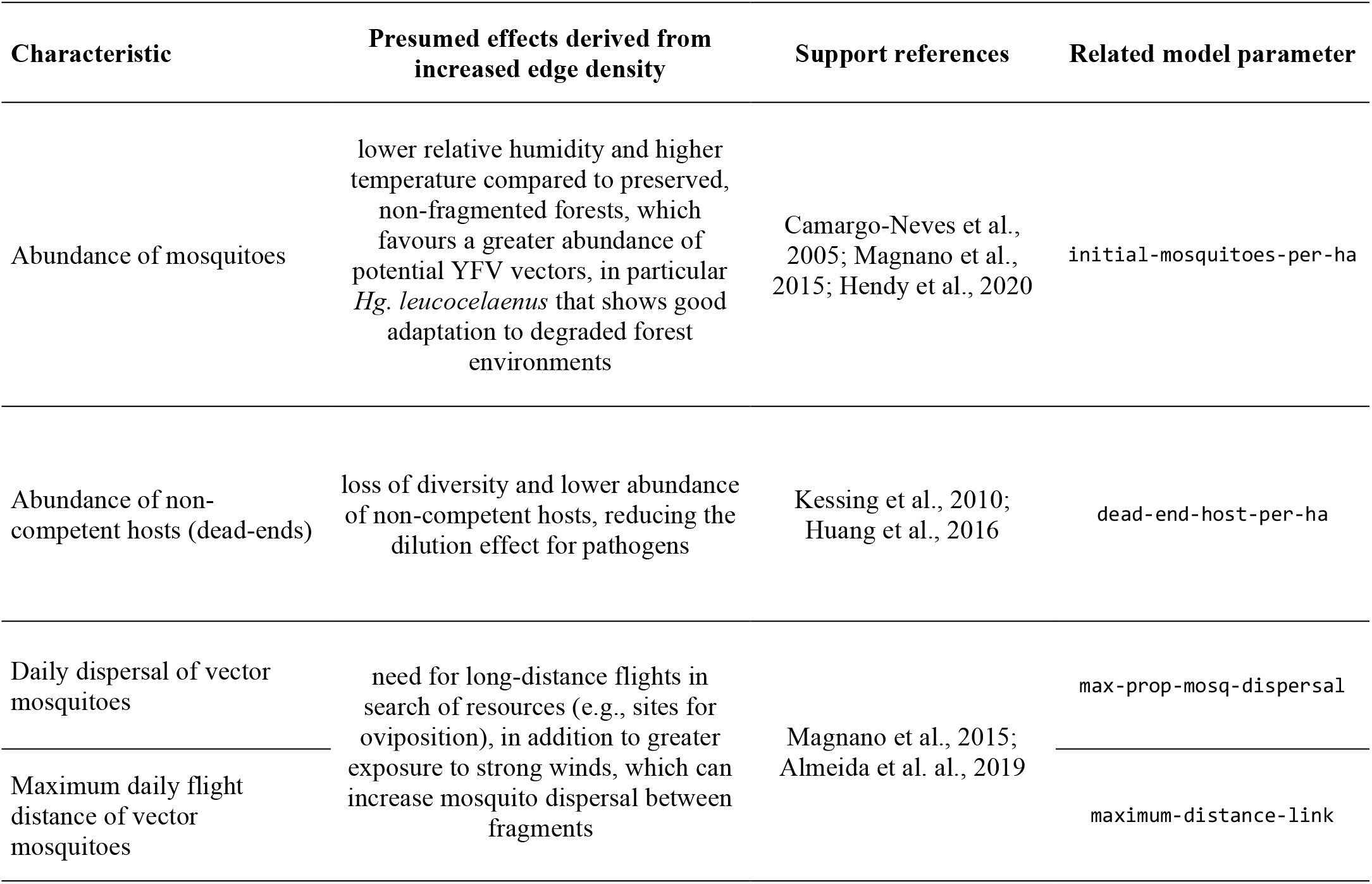
Assumptions made by the landscape-dependent scenario, including the characteristic, presumed effect, supporting references, and related parameter of the model.

| Characteristic | Presumed effects derived from increased edge density | Support references | Related model parameter |
| --- | --- | --- | --- |
| Abundance of mosquitoes | lower relative humidity and higher temperature compared to preserved, non-fragmented forests, which favours a greater abundance of potential YFV vectors, in particular *Hg. leucocelaenus* that shows good adaptation to degraded forest environments | Camargo-Neves et al., 2005; Magnano et al., 2015; Hendy et al., 2020 | initial-mosquitoes-per-ha |
| Abundance of non-competent hosts (dead-ends) | loss of diversity and lower abundance of non-competent hosts, reducing the dilution effect for pathogens | Kessing et al., 2010; Huang et al., 2016 | dead-end-host-per-ha |
| Daily dispersal of vector mosquitoes | need for long-distance flights in search of resources (e.g., sites for oviposition), in addition to greater exposure to strong winds, which can increase mosquito dispersal between fragments | Magnano et al., 2015; Almeida et al. al., 2019 | max-prop-mosq-dispersal |
| Maximum daily flight distance of vector mosquitoes |  |  | maximum-distance-link |

### 2.3. Simulation experiments

To simulate the transmission and propagation of YFV, six fragmented landscapes of the Atlantic Forest located in the state of São Paulo were selected (Figure 3) and represented using shapefiles obtained from the Forestry Inventory of the State of São Paulo 2020 (available at: https://www.infraestruturameioambiente.sp.gov.br/sifesp/inventario-florestal/). The areas in the State of São Paulo were selected due to the extensive surveillance and information system on yellow fever outbreaks in this State, made available by the Center for Epidemiological Surveillance of the São Paulo State Health Department (São Paulo, 2019). In all selected areas, there were epizootic outbreaks of YF with reported human cases between the period 2017 to 2018. The selected landscapes have distinct characteristics regarding the proportion of forest cover and edge density, selected to reflect the classification used by Ilacqua et al. (2021) to identify landscape fragmentation thresholds for YF. In this classification, the municipalities were divided into low (<30%), intermediate (30-70%), and high (>70%) forest cover and between those with equal or greater than 80 m/ha of edge density and those with less. For the present work, landscapes 1 and 2 present high forest cover (> 70%) and low edge density (< 80 m/ha). Landscapes 3 and 4 present low forest cover (< 30%) and high edge density (> 80 m/ha). In turn, landscapes 5 and 6 have intermediate forest cover (30 – 70%) and high edge density (> 80 m/ha) (Figure 3 and Supplementary information S2). The measurement of these landscape metrics was carried out in the Qgis v3.12.1 software (https://qgis.org) with the help of the Lecos plugin (Jung, 2013).

**Figure 3.**
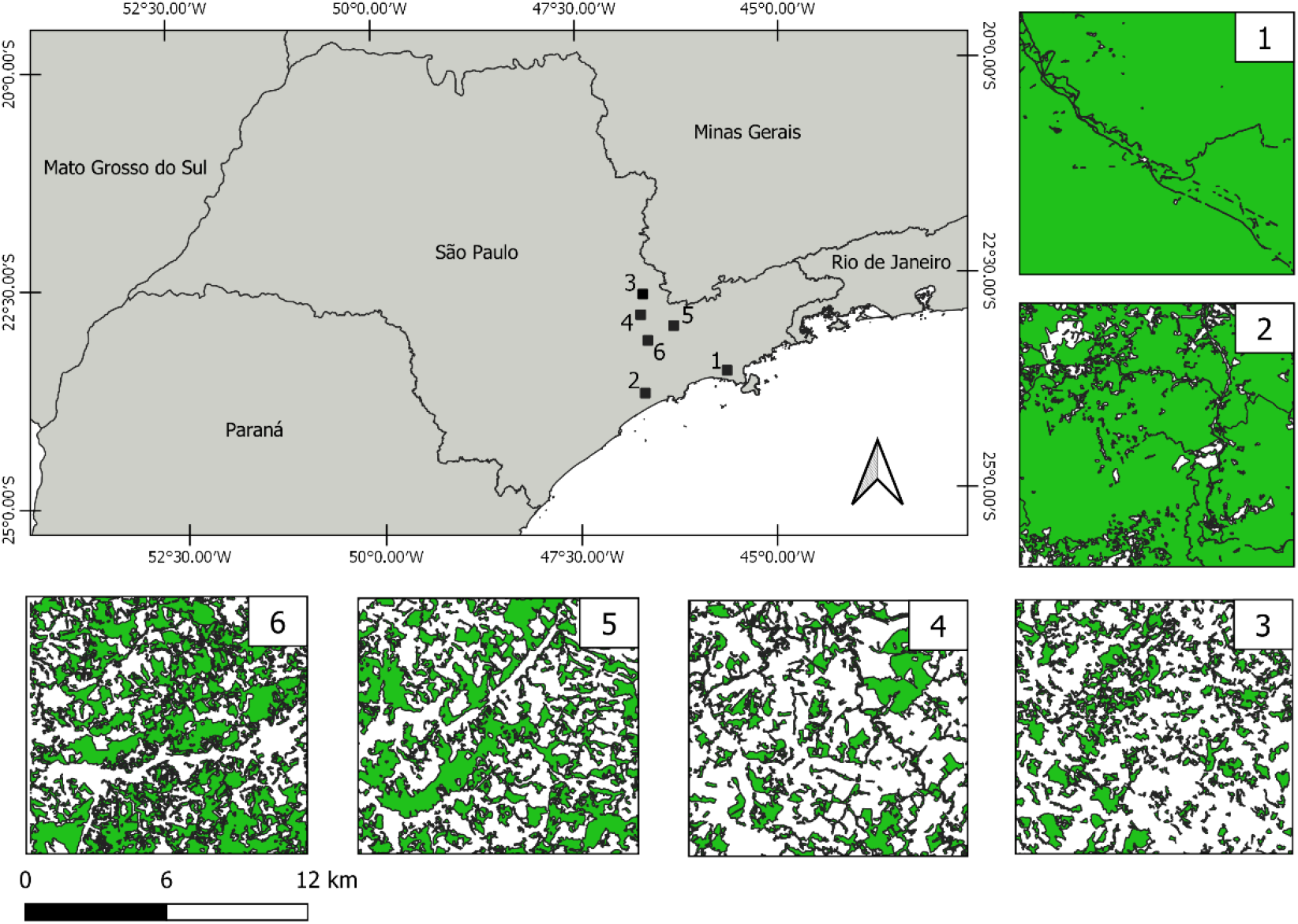
The location of six areas in the State of São Paulo used to simulate YFV propagation.

Two scenarios were assumed for the simulation, here named landscape-dependent and default. The landscape-dependent assumes that more fragmented landscapes favour higher permeability for viruses (Ilacqua et al., 2021; Priest et al., 2022; Wilk et al., 2022). In this scenario changes in edge density and forest cover influence the abundance and dispersal of the vector mosquitoes and the abundance of dead-end hosts in the landscape. In turn, the default scenario assumes that the spatial configuration of the landscape does not influence the abundance and dispersion of vector and dead-end hosts (null scenario). While in the default scenario, the only driver of variation in the virus propagation is the spatial configuration of the nodes and how they are spatially linked, in the landscape-dependent scenario is included a second driver of variation caused by changes in the abundance and dispersion of vectors and dead-end hosts, influenced by different combinations of edge density and forest cover. Table 4 provides information that supports the assumptions made by the landscape-dependent scenario.

Four parameters were tested for sensitivity to the two scenarios, as they are directly linked to the hypothesis of the effect of the landscape on the spread of the virus (as described in table 4). The parameters are 1) initial-mosquitoes-per-ha – which represents the initial and maximum seasonal abundance of mosquitoes per hectare, 2) dead-end-host-per-ha – representing the population size of dead-ends per hectare, 3) max-prop-mosq-dispersion - the maximum proportion of mosquitoes allowed to disperse to a linked node during a time step, 4) maximum-distance-link - the maximum distance in patches between two nodes to create a link, which influences the maximum daily distance that infected mosquitoes and hosts could disperse the virus. Values considered plausible were assigned to the four parameters and it was allowed that they could vary by up to 30%, thus limiting maximum and minimum values for the parameters. Thus, in the default scenario, the same values were attributed to these four parameters and tested for the three landscapes. In the landscape-dependent scenario, landscapes with more than 70% forest cover and less than 80 m/ha of edge density (landscapes 1 and 2) were tested with the maximum value for dead-end hosts (30% increase compared to default) and the minimum values for the other three parameters (a decrease of 30% compared to default), while landscape with less than 70% forest cover and more than 80 m/ha of edge density (landscape 3, 4, 5 and 6) were tested with the minimum value for dead-ends and maximum for the other three parameters. This information is summarized in Table 5.

**Table 5.**
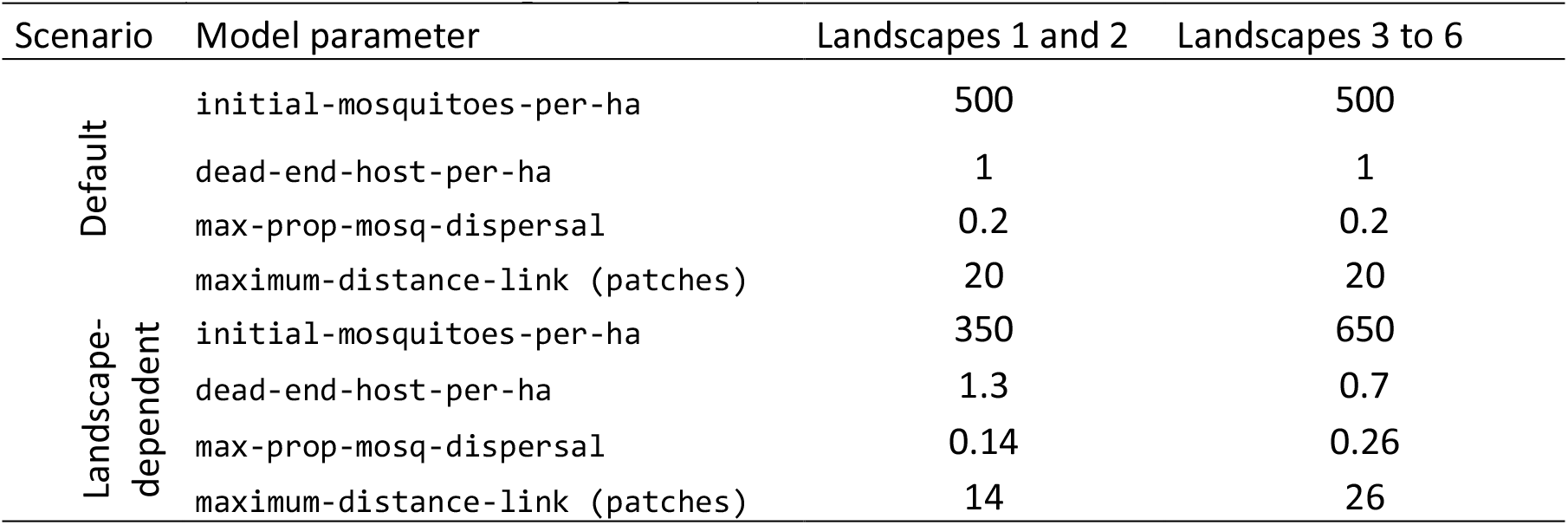
Assigned values to four model parameters according to two hypothetical scenarios (default and landscape-dependent).

After defining the scenarios and parameter initialization values, simulations were performed to assess whether the model outputs fit empirically observed patterns. For this purpose, seven patterns were evaluated according to data obtained in the literature (see Table 6). For each landscape and scenario, the parameter YFV-initial-burden was varied from 0.005 to 0.02 (intervals of 0.005), and the parameter start-YFV- circulation was set to four different values (183, 273, 365, and 455), representing the seasons (winter, spring, summer, and fall, respectively). For each set of parameters, five repetitions were performed totalling 960 runs.

**Table 6.**
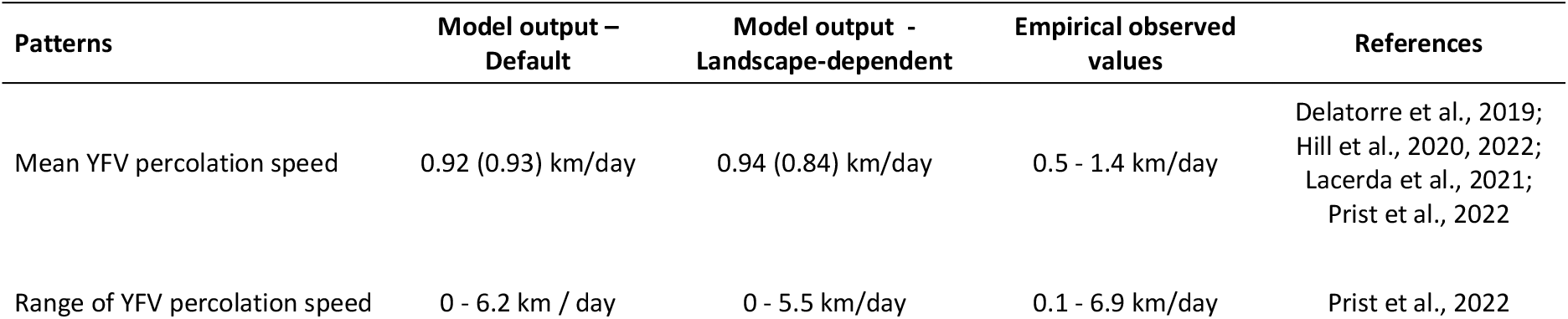
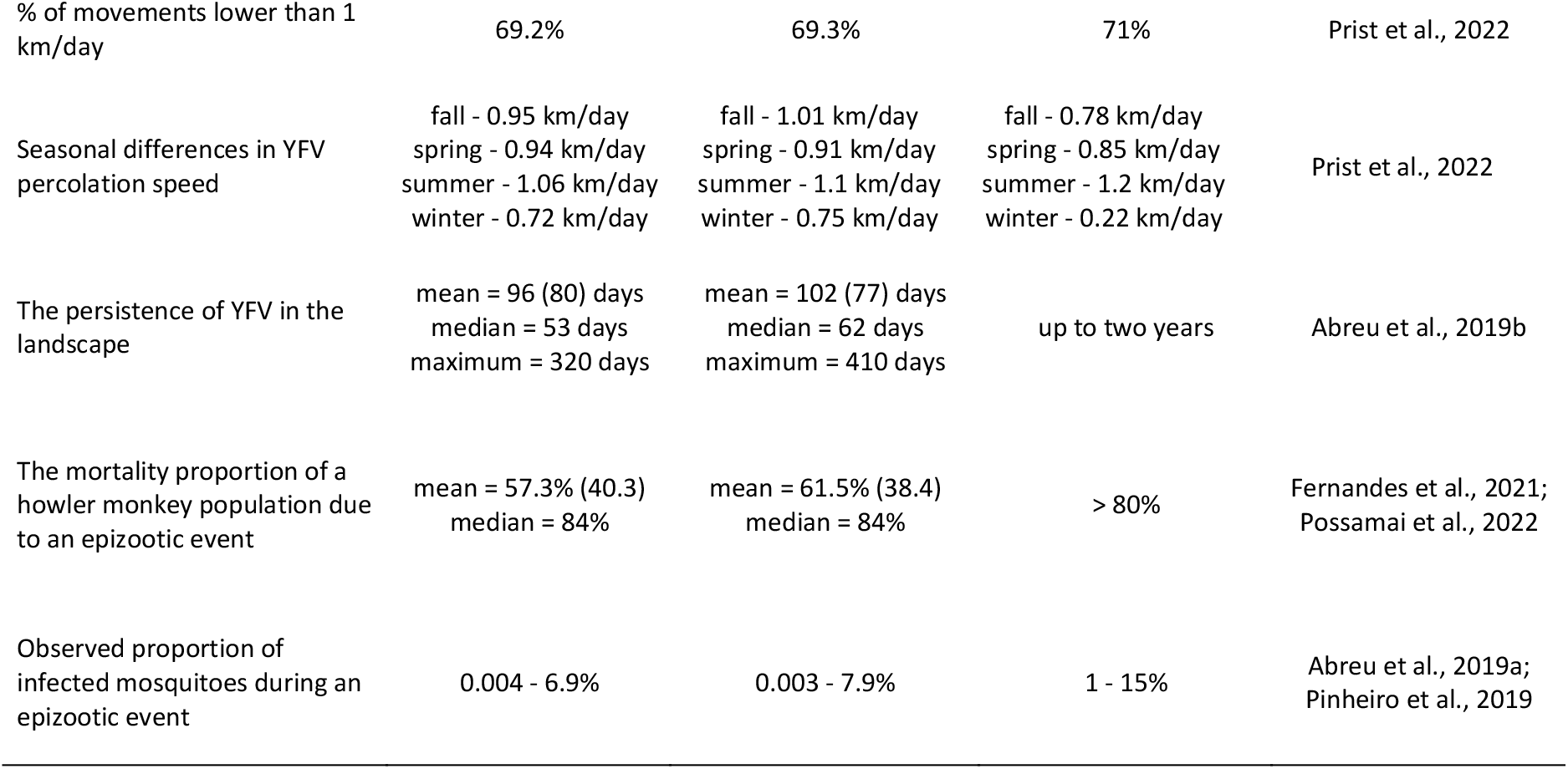
Values obtained by simulation of the model for seven patterns and comparison to observed empirical values. Two scenarios (default and landscape-dependent) were evaluated following different assumptions about the effect of landscape on YFV propagation. The values for each scenario were calculated based on simulations including the six landscapes. In parentheses are the standard deviations.

A second set of analyses was performed to evaluate the effect of different scenarios on two outputs: 1) percolation speed (km/day) – which calculates the virus propagation speed in the landscape and is measured as the distance of the initial area where the virus emerged (the centroid of this area) to the farthest node where the virus spread divided by the number of days. We opted to measure the average percolation speed for ten days after viral emergence, given that Prist et al (2022) observed that about 83% of YFV movements (time difference between two spatially related epizootics) occurred within this period after the virus detection at a source area; 2) spread proportion – which calculates the virus spread in the landscape, i.e., from the total number of receptive areas (nodes) to how many the virus has spread during the simulation, calculated as a proportion (number of nodes where the virus spread during the simulation divided by the total number of nodes). For each landscape and scenario, 100 repetitions were simulated, and the mean and standard deviation of the outputs were calculated. The parameters YFV- initial-burden and start-YFV-circulation were kept constant (0.02 and 365, respectively).

To evaluate the sensitivity of the outputs for each tested parameter and possible interaction effects, a global sensitivity analysis was performed using Sobol’s method (Sobol et al., 2007; Saltelli et al., 2010). The sample space of the input parameters was obtained using a Latin hypercube sample, and the values of each parameter were generated with three decimal digits using a uniform distribution. The variation in input parameters was limited to 30% around the default value. Because of the high computational cost of a Sobol sensitivity analysis, 500 samples were allowed for each input parameter, giving a total cost of 3000 model evaluations for each tested landscape, calculated as ’samples * (parameters + 2)’ according to the method. Confidence intervals for the estimates were obtained using 100 bootstraps. To ensure reproducibility of the analysis, the random number seed ’1’ was used.

The analyses were performed using the behaviour space tool available in NetLogo 6.1.1 (Wilensky and Shargel, 2002) and the ’nlrx’ package in the R computing environment (Salecker et al., 2019). The Sobol analyses were performed using the sobol2007 function available in the ’sensitivity’ package (Pujol et al., 2015). Given that even small effects can be found to be statistically significant after a sufficiently high number of replications (White et al., 2014), it was decided not to perform statistical tests to compare the effect of changes in input parameters on variations in the outputs. The model simulations were performed on a notebook with an Intel® Core™ i7 processor - 8GB RAM - and to optimize the time required for the Sobol analysis, the simulations were distributed on a HPC with 200 cores.

## 3. Results

The outputs obtained by simulations indicate that the model was able to reproduce empirical patterns in both hypothetical scenarios, presenting patterns that qualitatively and quantitatively approximate the available empirical data. Most outputs measuring different aspects of YFV presented values that were similar or within the observed ranges for the patterns (Table 6). The seasonal differences in YFV percolation speed showed higher values for the summer, intermediate in spring and autumn, and lower in winter, qualitatively similar to what was expected based on data. The main difference between simulation and empirical data was noted for the average speeds in winter, which were three to four-fold higher in the simulations compared to observed data (Table 6, Figure 4).

**Figure 4.**
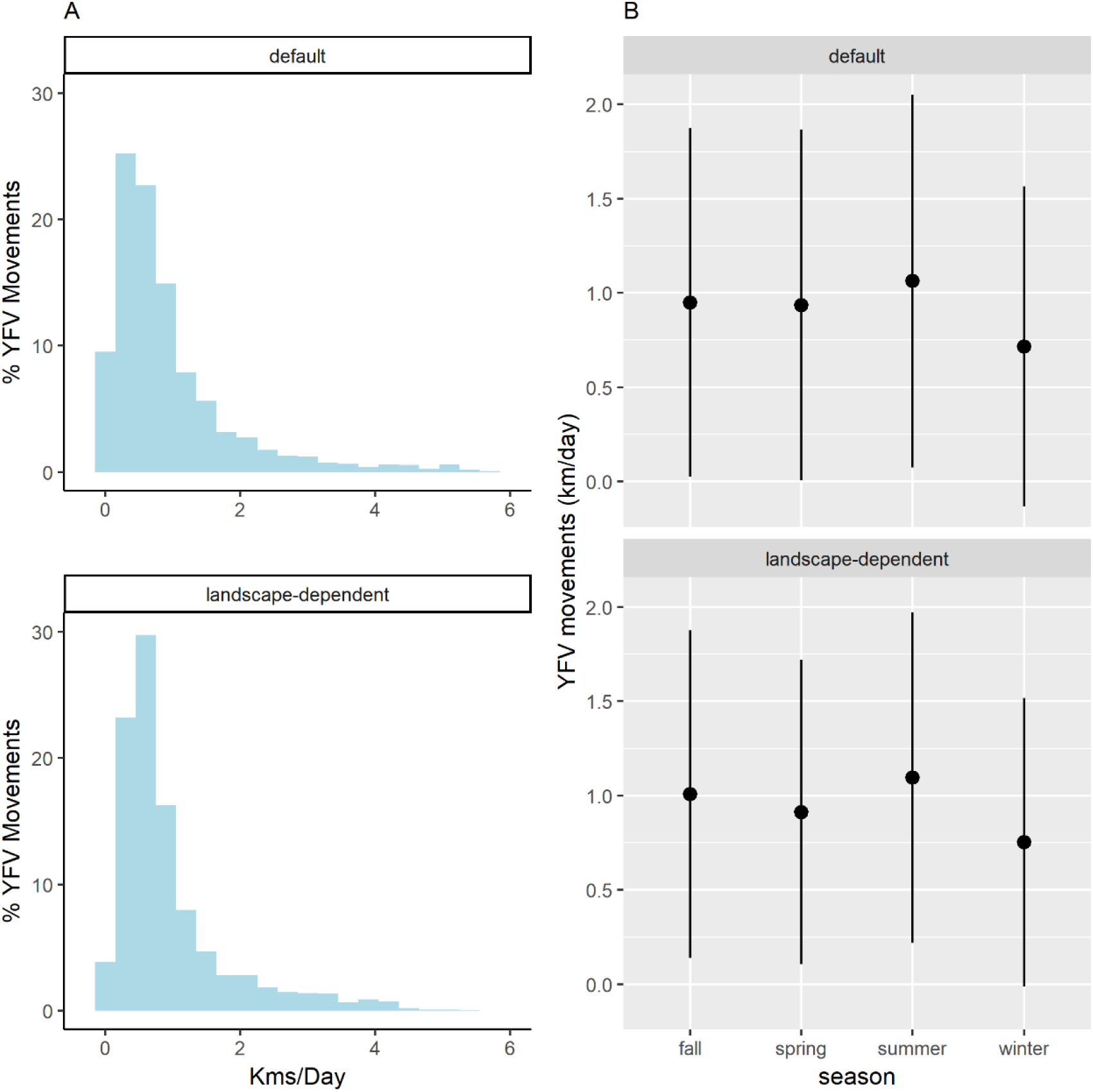
Outputs obtained through model simulations for (A) the total distribution of YFV movements and (B) the distribution of these movements by season. The values for each scenario (default and landscape-dependent) were calculated based on simulations including the six landscapes.

The simulations show that the speeds of the virus through the landscapes were different compared to the two hypothetical scenarios. In the default, the average speed was higher in the landscapes with a high proportion of forest cover and lower edge density, land 1 and 2 (Figure 5A). In the landscape-dependent scenario, landscapes with intermediate proportion of forest cover and high edge density, land 5 and 6, presented slightly higher average speeds than land 1 and 2. In turn, those with low forest cover and high border density, land 3 and 4, presented the lower average speed in both scenarios (Figure 5B). Differences between scenarios were also observed for spread proportion. In the default the average spread proportions were above 0.6 for land 1, 2, 5, and 6, achieving 100% for land 2, and below 0.35 for land 3 and 4 (Figure 5C). In the landscape-dependent scenario, land 1 and 2 presented a decrease in the spread proportion, while an increase was observed for the other landscapes. Except for land 2, all landscapes were above 0.8 in this scenario (Figure 5D).

**Figure 5.**
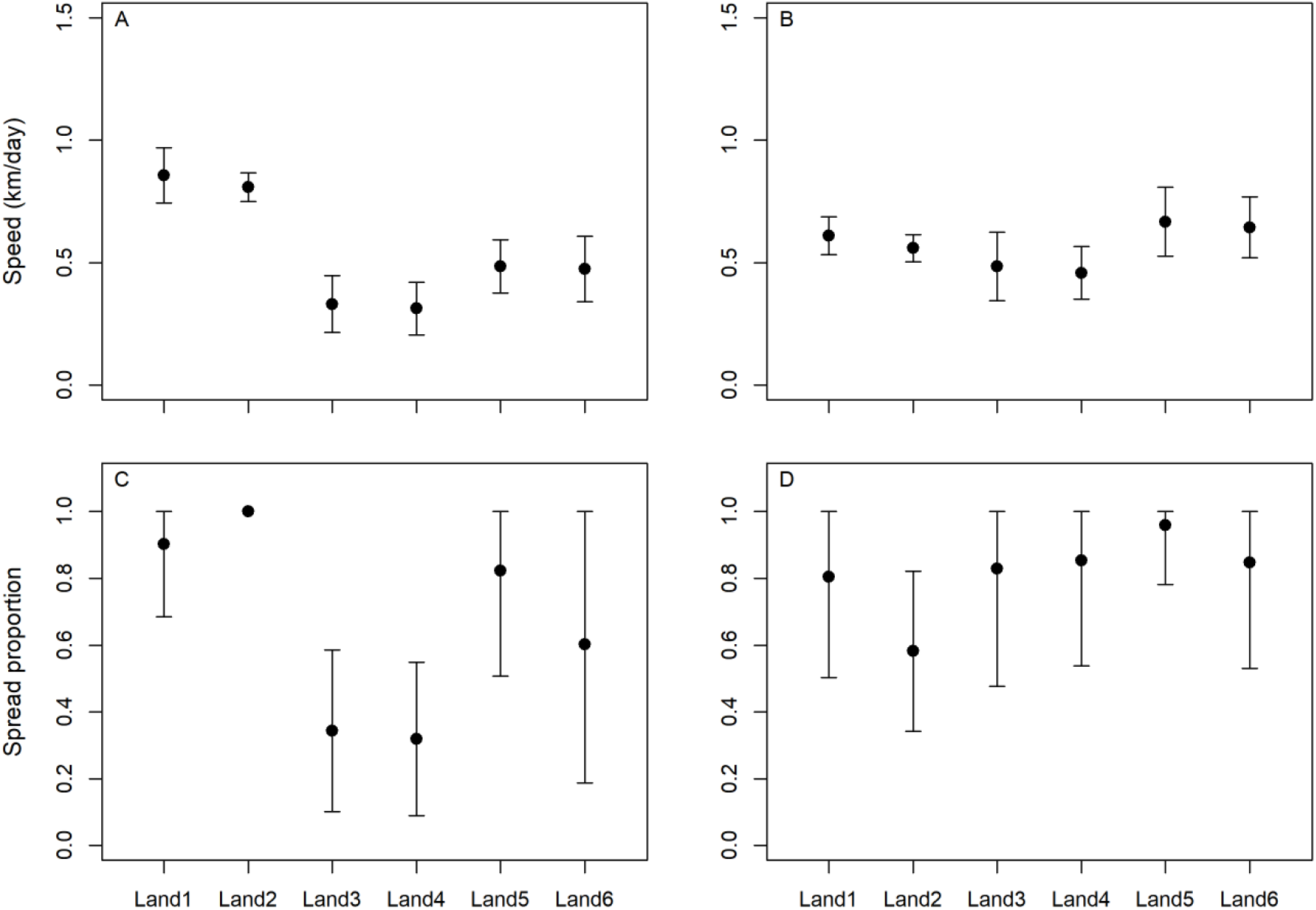
Model outputs for speed (km/day) and spread proportion of YFV for six landscapes (Land 1 to 6). In A and C, the parameters were set to default scenario, and in B and D the parameters were set to landscape-dependent scenario. Points represent the average values and error-bars the standard deviation. For each landscape and scenario, 100 repetitions were run.

The global sensitivity analyses indicate that the tested parameters have little or no main effect (Sobol first order index) on the outputs, but demonstrate relevant effects when considering the interaction between them (Sobol total index). Considering only the average total index, initial-mosquitoes-per-ha (mq) had in most cases superior values compared to other parameters (Figure 6).

**Figure 6.**
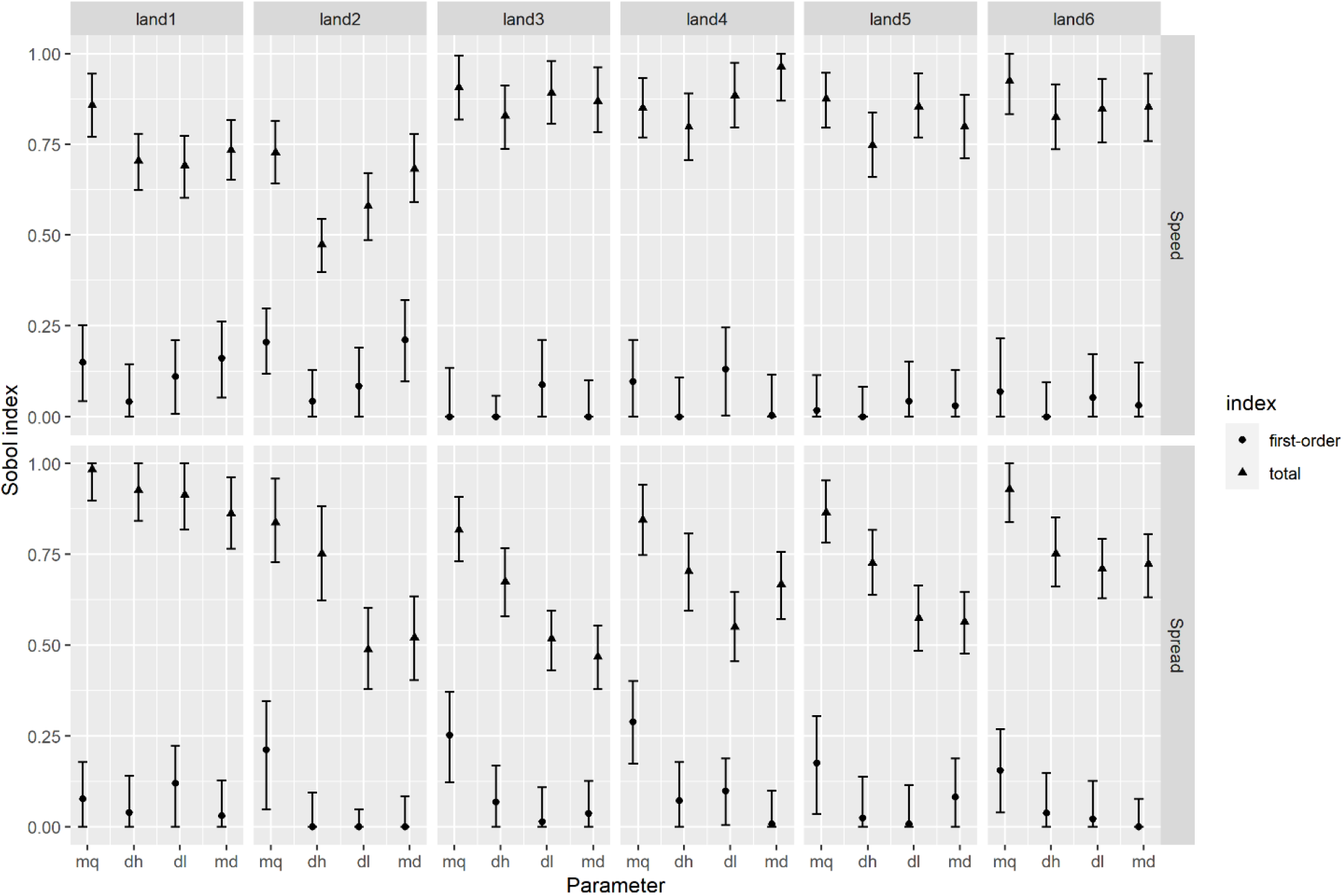
Sobol analyses present the first order and total index to four model parameters compared to two outputs (virus speed and spread) and six different landscapes. The parameters are initial-mosquitoes-per-ha (mq), dead-end-host-per-ha (dh), maximum-distance-link (dl), and max-prop-mosq-dispersal (md).

## 4. Discussion

### 4.1 Model applicability

We demonstrated that the developed model is able to dynamically simulate the spatial and temporal propagation of SYF in large fragmented landscapes. This offers new possibilities to investigate ecological and epidemiological aspects of SYF and can be used as a tool to support surveillance and prevention measures. Here, simulations were performed to investigate the hypothesis that fragmented areas facilitate a rapid spread of YFV. However, the model also has the potential to investigate other situations of interest in the ecology and epidemiology of SYF, both in theoretical and applied fields. Defining the corresponding scenarios and simulating them is facilitated by our detailed, complete model description, and by using NetLogo to implement the model, which is a free software platform and programming language that is easy to install and learn, runs on all major operation systems, and is well supported by its developers and its user community. These features made it, for example, possible that the complex honeybee colony model BEEHAVE (Becher et al., 2014), was used in more than 25 published studies, with more than half of them without any of the BEEHAVE developers involved.

In the theoretical field, the model enables exploration of circumstances wherein the virus could persist in endemic conditions, as observed in the Amazon region (Bryant et al., 2003). From identifying the conditions that result in endemic circulation, it will also be feasible to explore the factors that allow the necessary conditions for epizootic waves to reach extra-Amazonian regions (Possas et al., 2018). Another potential theoretical application (which can be extended to other vector-borne diseases) would be to investigate the impact of biodiversity on the maintenance and spread of YFV, since the model structure allows for the inclusion of host species with different population densities and levels of competence for the virus, thus allowing to test how the dilution effect operates in different configurations of communities and landscapes.

In an applied context, the model can be refined and calibrated to identify the most likely routes of virus spread and their speed of propagation, which is fundamental for the timely definition of priority areas for the prevention and immunisation of the population (Brazilian Ministry of Health, 2019). Future versions will simulate the presence of humans and test the level of immunisation required in the population considering landscape-oriented transmission dynamics to contain potential epidemics and support better management of vaccine stockpiles and material and human resources. In addition, the inclusion of urban vectors in the simulations would help identify situations that might favour the reurbanization of the virus (Couto-Lima et al., 2017).

Other vector-borne diseases, particularly those transmitted by mosquitoes in forest areas, have the potential to be investigated using the presented model. In this context, autochthonous malaria in the Atlantic Forest, transmitted by *Anopheles* mosquitoes of the subgenus *Kerteszia*, is an interesting example of a potential application of the model, since, similar to SYF, this *Plasmodium* transmission system involves sylvatic vector mosquitoes and NHPs as the main hosts (Laporta et al., 2013; Medeiros-Sousa et al., 2021). West Nile fever is another example of a disease with the potential to be simulated by the model since transmission dynamics include multiple wild vectors and host species (Chancey et al., 2015).

### 4.2. Reproduced empirical patterns

The two simulated scenarios were able to reproduce most of the tested empirical patterns, suggesting that both are plausible and provide a degree of realism for investigating the dynamics of SYF. The average viral percolation speed was close to 1 km/day, within the range of empirically observed values in which average speeds varied between 0.5 and 1.4 km/day (Delatorre et al., 2019; Hill et al., 2020; 2022; Lacerda et al., 2021; Prist et al., 2022). Regarding the range of speeds and percentage of movements lower than 1 km/day, the model outputs approximate the data observed by Prist et al. (2022). However, for seasonal differences in YFV speed, the simulated values were compatible with the observed values for summer, spring, and autumn, but were over three times higher in winter. It is expected that with the reduction in mosquito abundance during winter due to lower temperatures and low precipitation, YFV activity will decrease to low levels and even cease in many places (Hamlet et al., 2021). If this is a limitation of the model, future versions should include additional mechanisms and calibration to reproduce a slower virus percolation speed in the landscape during the winter period.

Whether YFV can persist in forest fragments for long periods after its emergence is a tricky question. Here, the outputs indicated average periods of about three months for YFV circulation in the simulated landscapes. Although it has been demonstrated that the virus was able to circulate in the same fragment for at least two years (Abreu et al., 2019b), what is generally observed is that the YFV tends to persist for short periods in forests outside its endemic area during outbreaks. The reason for this is that epizootic events usually cause high lethality in dense howler monkey populations harboured by these fragments, rapidly reducing susceptible hosts (Fernandes et al., 2021; Medeiros-Sousa et al., 2022).

With respect to the within-fragment patterns, the simulated proportion of howler monkeys that died was similar to that empirically observed when considering the median values of the outputs, with values above 80% in the two simulated scenarios. It is known that howler monkey mortality is quite high during YFV epizootics in the South American continent, which increases the risk of population decline in the coming decades (Moreno et al., 2015). During an epizootic that occurred between 2017 and 2018 in a forest fragment in Minas Gerais, Brazil, Possamai et al. (2022) reported an 86.6% decrease in the howler monkey population. Another good indicator of virus circulation intensity in the environment is the proportion of infected mosquitoes in a particular epizootic area. The simulated values show considerable variability, but with maximums not exceeding 10% of the population. For both *H. janthinomys* and *H. leucocelaenus*, the proportions of infected individuals in the population have shown considerable variation across the various field studies, with minimum infection rates ranging from 1% to over 15% of the sampled population. This variability may reflect the abundance of infected amplifying hosts and vector mosquitoes in the area (Abreu et al., 2019a; Pinheiro et al., 2019; Medeiros-Sousa et al., 2022).

### 4.3. The effect of fragmented landscape on SYF

We modelled the transmission dynamics of SYF in two scenarios to test the hypothesis that fragmented landscapes with high edge density enable a faster spread of YFV. The default scenario outputs do not support the hypothesis of facilitated spread in highly fragmented areas. This is because the virus exhibited higher average values of speed and spread in less fragmented areas with lower edge density. Conversely, in the landscape-dependent scenario, the simulations indicated that the areas with higher fragmentation, greater edge density, and intermediate forest cover exhibited faster average percolation speed. These findings are consistent with some recent studies (Prist et al., 2022; Wilk-da-Silva et al., 2022). However, it should be noted that even when significant differences in parameter values are considered, the divergence in mean output values between more preserved (landscapes 1 and 2) and less preserved (3 to 6) areas was small in this second scenario, both for speed and spread. This means that to assume that fragmented landscapes with high edge density facilitate the spread of YFV, there must be significantly different conditions between them and more preserved areas, at least regarding the abundance and dispersion of vectors and dead-end hosts.

The recent studies that examined the spatial and temporal distribution of virus spread using reported outbreaks in the state of São Paulo between 2016 and 2018 support the hypothesis that fragmented areas can facilitate the rapid spread of YFV. Using reported epizootics, Prist et al. (2022) carried out analyses based on network and circuit theory to identify the landscape features that enable or hinder virus connectivity. The study examined multiple models, and the most favourable one indicates that the spread of YFV occurs primarily through roads adjacent to forested areas and along forest edges at the interface with agriculture and water. In a similar analysis, Wilk-da-Silva et al. (2022) used mixed-effects models to investigate whether models that include only the effect of geographic distance on virus spread are improved by variables that measure YFV permeability across the landscape associated with different land uses. The best-fitting model indicates higher permeability associated with geographical distance and forest edges at the interface with water, agriculture, non-forest formation, and forestry. Additionally, both studies cited indicate that urban and forested core areas are important barriers to virus spread.

The potential impact of forest fragmentation on the increased permeability for YFV may be attributed to disturbances to the vector and host populations caused by the loss and alteration of their habitat characteristics. Studies indicate that *H. leucocelaenus* tends to be more abundant in areas of degraded forest, as this species adapts to these conditions. Added to this is the high dispersal capacity of *Haemagogus* mosquitoes, capable of flying up to 11 km in search of blood sources and breeding sites to lay eggs. The dispersal can also be passive, as the mosquitoes are carried by the wind between fragments separated by open areas or roads (Causey et al, 1950; Camargo-Neves et al., 2005; Magnano et al., 2015; Almeida et al., 2019; Hendy et al., 2020). Furthermore, habitat loss and fragmentation in areas with high biodiversity may result in a decrease in species abundance or local extinction of species that act as barriers to virus transmission due to their lower competence as hosts. In this way, the dilution effect of the pathogen would be less effective in areas with higher edge density due to fragmentation, compared to more continuous and well-preserved forest areas that provide greater diversity of non-competent hosts (Kessing et al., 2010; Huang et al., 2016).

On the other hand, observations suggesting facilitated spread of YFV in more fragmented areas and along forest edges may be subject to bias because they are based on the locations and dates of encounters of dead NHP, and most reports of epizootics are based on the finding of their carcasses in the edge of forests, at the interface with roads, in rural areas and around urbanised areas, where access and visualisation of events are easier than in core areas of continuous and less fragmented forests (Fernandes et al., 2021). In addition, the reports in preserved forests are subject to delays, as the virus may have been circulating in a given site for weeks or months without being noticed before. Thus, additional studies are needed to clarify whether what is observed is truly a relationship between edge areas and SYF dynamics or simply results from surveillance limitations of epizootics.

As observed in the simulations, landscapes with a lower proportion of forest cover (3 and 4) showed a slower average speed for virus percolation compared to other landscapes tested. Also, landscapes with lower forest cover showed a reduced capacity for virus spread compared to others when the default scenario was simulated. This corroborates with a theoretical model developed for Atlantic Forest that has identified a critical threshold of remaining forest cover whose capacity for ecological resilience is compromised, causing a regime shift in biodiversity that can affect zoonotic diseases through changes in host-vector-pathogen interactions. Such thresholds are around 30%, below which connectivity between remaining fragments tends to be severely compromised (Pardini et al., 2010). Ilacqua et al. (2021) adopted this threshold as a metric for assessing the risk of YFV occurrence in 151 municipalities in central and southeastern Brazil where epizootics and human cases were reported between 2014 and 2019. The investigation identified that municipalities with 30-70% forest cover and >80 m/ha edge density had an 85% increased relative risk of YFV occurrence compared to municipalities with >70% or <30% forest cover.

### 4.4. Global sensitivity analysis

Global sensitivity analyses consistently showed, for the different simulated landscapes, that the outputs were more influenced by the interaction between the tested parameters than by individual effects. This suggests that a combination of factors acting synergistically determines the speed and magnitude of YFV epizootic and epidemic waves. Among the parameters tested, the total Sobol index was slightly higher for mosquito abundance compared to other parameters. Likewise, simulations conducted by Medeiros-Sousa et al. (2022) indicated that mosquito abundance is also an important parameter for influencing infection prevalence in mosquitoes, primate mortality, and YFV persistence within a fragment, especially when interacting with other key parameters. For YFV, previous studies have shown that periods of climatic anomalies that cause changes in rainfall and temperature increase the likelihood of large YF epizootics and epidemic waves when mosquito numbers exceed normal levels (Vasconcelos et al., 2001; Vasconcelos, 2010; Zhao et al., 2018). In addition, factors that influence mosquito densities such as rainfall, temperature, humidity, and environmental imbalances in small forest patches have been linked to YF epizootics and human infections (Almeida et al., 2019; Abreu et al., 2022).

### 4.5 Model limitations

The ability of the model to reproduce empirically observed patterns has been demonstrated, but important limitations must be considered in future versions of the model. The model only considers the structural connections between fragments, without considering how functional are these dispersal corridors for the virus. This is because the landscapes used in the simulation were classified in a simplified manner into forested or non-forested areas, without considering other potential barriers to the dispersal of vectors and hosts, such as high altitudes, urban areas or environments with low potential for the dispersal of vectors and hosts (*e. g.* monocultures of *Pinus* spp.) (Wilk-da-Silva et al., 2022). It is worth pointing out that the calibration and validation were largely based on data obtained in dense ombrophilous forest areas of the Atlantic Forest, and therefore it is not certain that the same patterns would be observed when considering landscapes of other vegetation types where YFV circulates, especially those of the Cerrado and Amazon biomes.

Many model parameters were assumed based on expert knowledge or considered within plausible values due to the absence of empirical investigations to define the value of such parameters, which increases uncertainty regarding the accuracy of the outputs. Furthermore, in the current version, the model does not incorporate human beings or urban vectors in the transmission dynamics, making it impossible to investigate some important aspects of YF epidemiology, such as the minimum level of population immunisation required to prevent epidemics and the risk of virus reurbanisation.

## 5. Conclusions

A hybrid compartment and network-based model structure was presented, capable of dynamically simulating the transmission and spread of SYF in fragmented areas. This allows the simulation of epidemiological, reproductive, and dispersal dynamics for a large number of individuals of different species on a broad landscape scale. The model was able to reproduce most of the tested empirical patterns and simulations suggest that under certain conditions, the virus may spread more quickly in fragmented environments with an intermediate degree of forest cover compared to either well-preserved or highly degraded areas. It is expected that the model will be applied to investigate various other features of SYF and other vector-borne diseases, thereby contributing to a better understanding of the ecology and epidemiology of these diseases and supporting surveillance, prevention, and control actions.

## Supporting information

Supplementary Information S1 - ODD protocol

## Acknowledgments

We would like to thank our colleagues at the Helmholtz Centre for Environmental Research for their contributions and support during the development of the work, especially Dr Karin Frank and Dr Jürgen Groeneveld. We also thank Renato Pereira de Souza of the Adolf Lutz Institute for valuable comments during the early stages of model development.

