## Supplementary Information S1 - ODD protocol for "Modelling the transmission and spread of yellow fever in forest landscapes with different spatial configurations"

#### ODD protocol

This model description follows the ODD (Overview, Design concepts, Details) protocol for describing agent-based models (Grimm et al. 2006, 2020). The model was implemented in NetLogo (Wilensky 1999), version 6.3.0 and is available at https://github.com/aralphms/YFVLandModel

##### 1. Purpose and Patterns

The model is designed to represent the transmission dynamics and propagation of the yellow fever virus (YFV) across fragmented landscapes and is based on the local agent-based model by Medeiros-Sousa et al. (2022). The emergence and transmission of sylvatic yellow fever (SYF) between wild mosquitoes and monkeys, and its dispersion between forest fragments is simulated. This simulated process allows exploring how different characteristics of vector, host, and landscape influence the spatial and temporal dynamics of the virus. The main question addressed is: Do highly fragmented landscapes facilitate the spread of YFV during epizootic waves? The purpose of the model is to better understand and predict spatiotemporal dynamics of YFV infections by (1) testing scenarios where the abundance of mosquitoes and dead-ends are dependent on both forest cover and edge density (see below), and (2) contrasting different spatial configurations of forest fragments.

Recent empirical observations from epizootic waves that occurred in Southeast Brazil between 2017 – 2019 have suggested that highly fragmented forested areas are more permeable to YFV (Prist et al., 2022, Wilk-da-Silva et al., 2022) and increase the risk of SYF disease occurrence in monkeys and humans (Ilacqua et al., 2021). The main hypothesis is that edge areas of these forests have higher abundances of the main vector, mosquitoes, and less diversity of dead-end hosts, which increases the contact rates between vectors and monkeys and facilitates virus amplification and propagation in the landscape. In addition, fragmented areas may facilitate the dispersal of mosquitoes for long distances looking for blood sources and breeding sites or being carried by the wind. Thus, high permeability to the virus is expected in highly fragmented landscapes with an intermediate degree of forest cover since these areas tend to present high edge density, support large populations of monkeys and mosquitoes, have a lower diversity of dead-end hosts, and have good connectivity between fragments allowing higher dispersal of mosquitoes and hosts.

The metrics and criteria used to claim that the model is realistic enough for its purpose are the following empirically observed patterns. Some of the individual patterns might be easy to reproduce and thus not necessarily indicate high realism of the model. However, making the model reproduce them simultaneously is challenging but has higher chances to capture the mechanisms underlying the spread of YFV in fragmented landscapes with sufficient realism ("pattern-oriented modelling", Grimm et al. 2005, Grimm and Railsback 2012).

1) **The average and variation in the YFV spreading speed** – different studies suggest that the virus spreads about 1km (0.6 – 1.4) per day on average in fragmented landscapes of southeastern Brazil (Hill et al., 2020; Lacerda et al., 2021; Prist et al., 2022). Movements from 0.1 to 6.9 km/day were observed, and from these 71% were up to 1 km/day (Prist et al., 2022).

2) **The seasonal differences in YFV spreading speed** – the average spreading speed turned out to be higher in summer, similar in fall and spring, and lower in winter (Prist et al., 2022).

3) **The persistence of the virus in the landscape** – In South America, YFV has a periodic and transitory characteristic outside its endemic area (Amazon region), propagating in an epizootic-epidemic wave that circulates in the forests until the stock of susceptible hosts is exhausted or the virus finds environmental restrictions to remain circulating (Possas et al., 2018; Almeida et al., 2019). Although YFV was observed circulating for up to two years in the same Atlantic Forest fragment (Abreu et al., 2019A), to date there is no evidence of an enzootic-endemic establishment of the virus outside the Amazon region.

In addition, local patterns observed within a forest fragment can be also used as criteria for model usefulness. Here, we consider as useful patterns the high mortality of the main host, expected to be >80% for howler monkeys (Fernandes et al., 2021; Possamai et al., 2022), and the observed proportion of infected mosquitoes during the peak of epizootic events (~1 – 15%) (Abreu et al., 2019B; Pinheiro et al., 2019).

#### 2. Entities, state variables, and scales

For a better comprehension of the model structure, this section is divided into three subsections. In subsection 2.1 we present the model scales, in 2.2 we describe the entities and their state variables, and in 2.3 the rationale and implementation of the compartment-based structure are described.

##### 2.1. Scales

The model is spatially explicit and uses either hypothetical configurations for model testing and analysis or shapefiles of real landscapes to simulate YFV propagation. The landscape consists of 121 x 121 patches, representing an area of approximately 144 km². The boundaries are assumed to be closed, i.e., there is not interaction with the world outside the model landscape. A shapefile of 144 grid cells representing 1 km² each is superimposed to the landscape to form the landscape units. The time step of the model represents one day, and the simulation comprises the period from the emergence of the virus to its disappearance in the landscape or when the simulation reaches 1825 days (5 years of 365 days).

##### 2.2. Entities and state variables.

Model entities include patches, forest fragments, landscape units, nodes, and links (Figure 1). Patches are the discrete spatial units of the grid defining the modelled landscape and can be of two types: forest or non-forest. They represent one hectare each. All patches have state variables for their coordinate on the grid, the ID of their landscape unit, and if they pertain to a initiator node (nodes that start the virus circulation during model simulation). Forest patches have additional state variables for the ID of their forest fragment and the ID of the node they belong to (see below). Forest *fragments* are formed by a set of patches of the type 'forest'. Landscape units are defined by a grid of larger spatial units of 1 km² each. A landscape unit can contain none, one or multiple forest fragments. Large forest fragments can be located in two or more landscape units. Both forest fragments and landscape units have ID numbers as state variables. The state variables of the patches, forest fragments, and landscape units are constant, as they provide, during initialization, the spatial information for the nodes (Table 1).

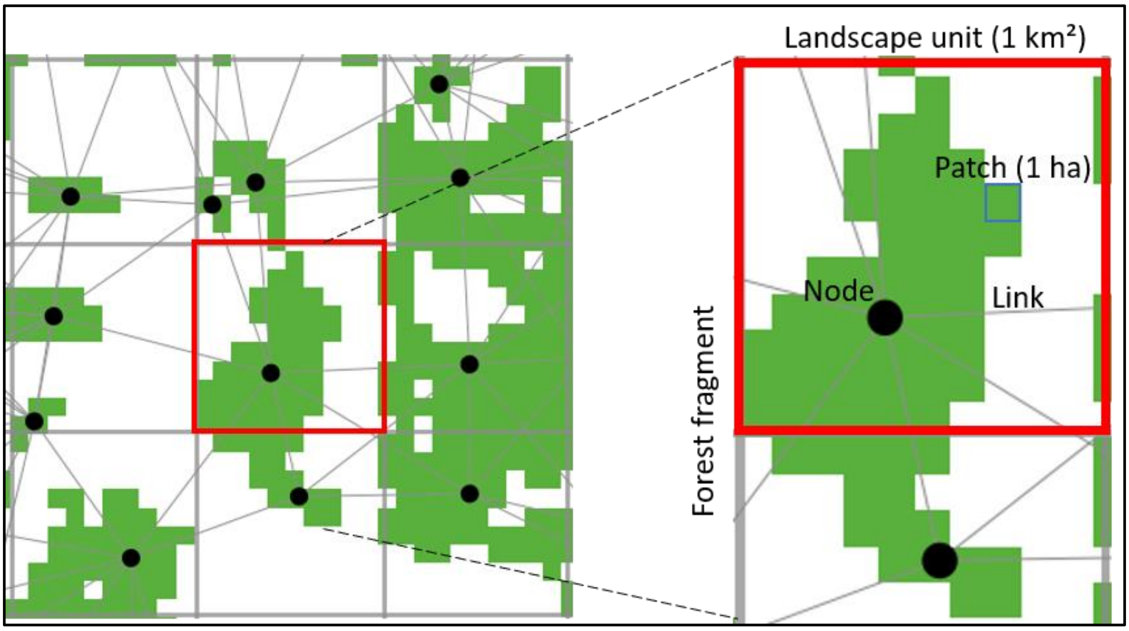


**Figure 1**. Example landscape to illustrate the model entities. The zoomed forest fragment has part of its patches located in one landscape unit (red square) and another part located in a second landscape unit. Thus, the model assigns two nodes (black circles) for the fragment. Each node has links to its neighbors.

Table 1. Description of the state variables assigned to each entity of the model. The structure and rationale of the four lists characterizing a node are discussed below (Section 2.2).

| Entity | State variable name | Description |
|---|---|---|
| forest fragment | ID | the number that identifies the fragment |
| landscape unit | ID | the number that identifies the landscape unit |
| patch | coordinates | the coordinates x and y in the landscape |

| | | |
|---|---|---|
| | patch-type | if the patch is forest or non-forest |
| | fragment-id | the id of the fragment where the patch is located |
| | landscape-unit-id | the id of the landscape unit where the patch is located |
| | node-id | the id of the node that represents the area where the patch is located |
| | patch-init? | if the patch pertain to a initiator node |
| node | coordinates | the coordinates x and y in the landscape |
| | my-node-id | the id of the node |
| | my-fragment-id | the fragment number of the patches represented by the node |
| | my-landscape-unit-id | the id of the landscape unit where the node is located |
| | my-grid-size | the number of patches the node is representing |
| | my-patches | agentset containing the patches that belong to the node |
| | mosq-list | list containing the number of mosquitoes in different reproductive and epidemic states |
| | host-list | list containing the number of hosts of different types and their epidemic states |
| | total-immatures | list containing the total number of immatures |
| | pot-infec-immatures | list containing potentially infected individuals from the total-immatures list |
| | new-eggs | the total number of eggs laid by gravid mosquitoes in a single day |
| | new-pot-infe-eggs | the total number of new potentially infected eggs |
| | active-transmission? | indicates whether the virus is circulating in the area |
| | external-node? | if the node is external or internal to the landscape |
| | prob-find-host | the daily probability of a blood-seeking mosquito finding a host |
| | viral-introduction | number of new arrivals of infected mosquitoes or hosts from other nodes |
| | initiator? | if the node initiates the virus circulation in the landscape |
| | current-k | the current carrying capacity for the mosquito population in the node |
| | k-max | the initial mosquito abundance and maximum value of current-k |
| links | minimum-distance | the minimum distance between the two represented areas, calculated based on the closest patches |
| | nodes-distance | the distance between the nodes of the link |
| | dispersion-rate-end1 | the proportion of mosquitoes that can disperse from end1 to end2 |
| | dispersion-rate-end2 | the proportion of mosquitoes that can disperse from end2 to end1 |
| | host-movement-rate-end1 | the proportion of hosts that can move from end1 to end2 |
| | host-movement-rate-end2 | the proportion of hosts that can move from end2 to end1 |

Nodes are abstract immobile entities that represent, for a forest fragment, the different reproductive and epidemic states of both mosquitoes and vertebrate hosts. Nodes were introduced to avoid a spatially explicit representation of vector and host dynamics within forest fragments, which would have led to too high computation times. To avoid artefacts of this spatially implicit representation of local dynamics, landscape units are used to make sure that large fragments are represented by more than one node. Thus, each forest fragment or part of a forest fragment within a landscape unit will be represented by a node (Figure 1). Each node has state variables for its position on the grid (the geometric center of the set of patches it represents), the IDs of the forest fragment and landscape unit they belong to, if the node is external or internal (external nodes are in the boundary of model world area and are responsible to initiate the virus propagation), the number of forest patches it represents, and if viral transmission is taking place at the area. Each node contains four different lists that represent mosquitoes (one list for adults and two for immatures) and host populations (one list that includes monkeys, alternative hosts, and dead-end hosts). The lists items (hereafter compartments) indicate the number of individuals in the different reproductive and epidemic states in the population (see detailed explanation in the Submodels section). Links indicate which nodes are spatially linked in the grid and determine the exchange of vectors and hosts between nodes (see sections 5, 7.10 and 7.11). The state variables of links include the calculated dispersion rates for mosquitoes and hosts between the two nodes, the minimum distance between the two represented areas, and the distance between the nodes (Table 1).

##### 2.3. Compartment-based model structure

The compartment-based structure of the model is a faster and computationally viable alternative to the agent-based model developed by Medeiros-Sousa et al. (2022). It allows us to simulate the transmission dynamics in fragmented landscapes with thousands or even millions of mosquitoes. In this compartment-based version, instead of individual agents representing mosquitoes or hosts, different compartments represent the number of individuals in a given state or combination of states and for a given node. The numbers in the compartments are integers. When a function of a submodel generates a floating point this value is rounded to the closest integer. Three compartment-based structures are used in the model, one for adult mosquitoes, a second for immature forms, and a third for hosts. Matrices of $m \times n$ states can be used to represent the compartments and transitions, as illustrated in figure 2.

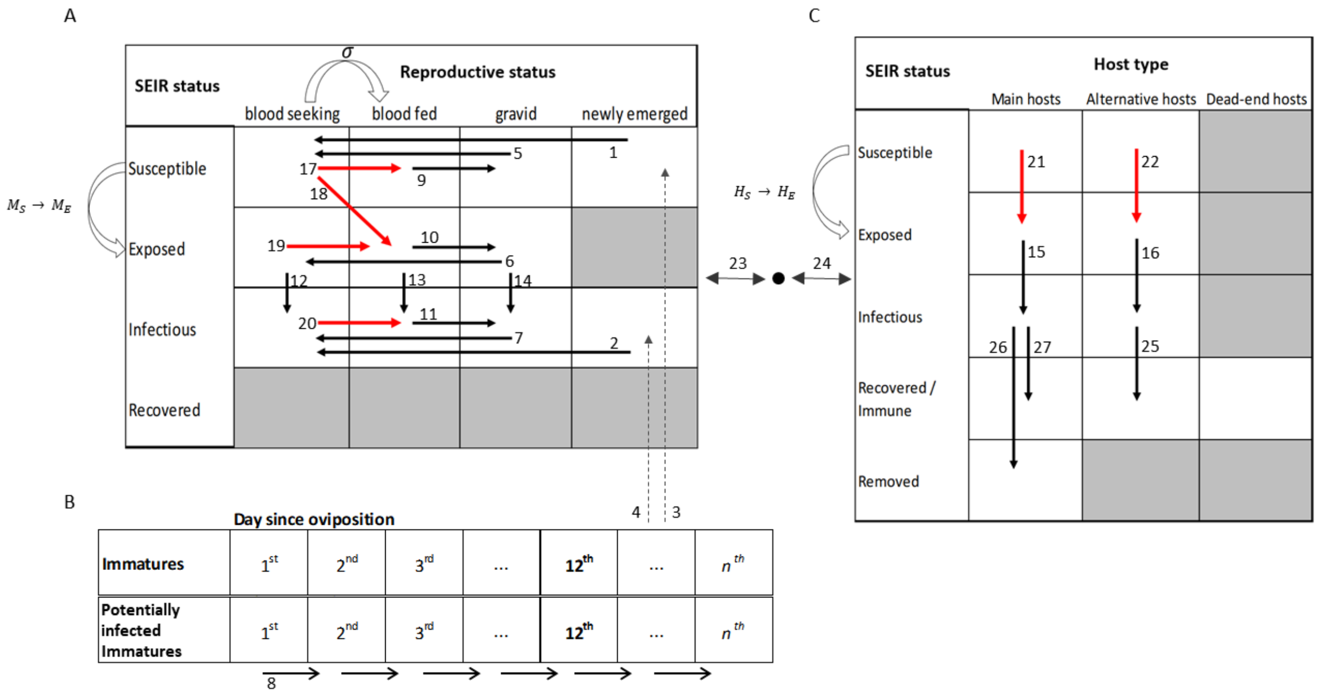


**Figure 2**: Representation of the compartment-based structure of the model. A - compartments for adult mosquitoes showing possible combinations (white compartments) between SEI and reproductive status. B – Compartments for immature forms representing the number of individuals at sequential different days after ovipositions. C – Compartments for hosts showing SEIR states. The arrows represent the possible compartment transitions and the numbers represent the order in which the transitions occur in the model (the same numbers are used, e.g. "Transition 1", as comments in the NetLogo program so that the corresponding code can easily be found). Red arrows indicate transitions that require host-vector interactions (σ) and change the state of mosquitoes from blood-seeking to blood-feeding (curved arrow on top of A). Infective interactions (curved arrows to the left sides of A and C) make susceptible individuals change to the exposed compartment ($M_S \rightarrow M_E$ or $H_S \rightarrow H_E$). Dashed lines represent the transference of individuals from the immature to adult compartments. Double arrows represent the transference of individuals (coming from different compartments) from and to a neighbor node (black circle).

Different compartments represent the circulation of the virus for mosquitoes and vertebrate hosts following an SEIR framework (Mandal et al., 2011): susceptible, exposed (the individual has been infected but is not yet able to transmit the virus during a latent period), infectious (the individual can transmit the virus) and recovered or removed. The compartment-based structure for mosquitoes can be represented by three states for viral infection (susceptible, exposed, and infectious), represented by rows, and four reproductive states (blood-seeking, blood-fed, gravid, and newly-emerged), represented by columns. In this matrix, the compartments for recovered and the combination exposed and newly emerged are considered impossible states in the population dynamics (grayed out), as it is assumed that these states do not occur in nature. In total there are 11 states (blank compartments) and 15 possible transitions (arrows) (Figure 2A).

The immature forms are represented by sequential compartments that form the breeding site structure in the model. Each compartment indicates the number of individuals and represents the time in days since these individuals were ‘laid’ at the breeding site. For example, the first compartment will hold individuals that were oviposited one day ago (the previous time step), the second will hold those laid two days ago, and so on. Once the individuals reach the minimum development time, the 12th compartment, they are allowed to move to adult newly-emerged compartments, but the number of days taken from the egg to the adult is variable depending on the conditions of the model environment (Figure 2B). In addition, a second list is used to count the number of potentially infected individuals in each compartment of the immature list. The individuals accounted in this second list represent the progeny of infectious mosquitoes and therefore they have a probability (determined by the `vertical-infection-rate` parameter) of having been infected via transovarial transmission (see details in sections 7.2, 7.5, and 7.6).

Host populations can be represented by SEIR status in the rows and by different types of hosts in the columns. Three types of hosts are considered: 1) main hosts – representing howler monkeys that are highly susceptible and have high lethality when infected with YFV; 2) alternative hosts – species that can become infected and transmit the virus to mosquitoes but have little or no lethality; 3) dead-end hosts – species that can become infected and even show symptoms but do not produce enough viremia in the bloodstream to infect new mosquitoes, becoming a barrier to the amplification of the virus. Given that individuals can recover from the infection and become immune or they can die and be eliminated from the simulation, the letter R of SEIR means 'Recovered/ Immune' or 'Removed' and can be represented by two different compartments. The dead-end hosts were allocated only in the Recovered/Immune compartment as these hosts cannot transmit the virus (although they may present all SEIR states in nature). Furthermore, the Removed compartment is disregarded for alternative hosts and dead-ends as for model simplicity it is assumed that these hosts do not die from the YFV infection and have constant population size (Figure 2C).

Each node (Table 1) has all the compartments. The transitions between compartments are updated at each time step and the processes used for these transitions will be explained in more detail in section 7.

#### 3. Process overview and scheduling

The model uses both a compartment-based and an network approach. Most of the procedures in a time step are executed by the nodes and represent the local dynamics by promoting changes in the mosquitoes and host compartments (section 2.2). These compartment-based procedures include those applied to population dynamics (reproductive status and mortality) and those for transmission dynamics (susceptible, exposed, infectious and immune compartments). The network structure of the model is limited to the exchange of values between linked nodes, which represents the movement and dispersal of individuals. Table 2 shows the order of events performed at each time step and gives a summarized description. Each procedure is a submodel that is detailed in section 7.

**Table 2**. Procedures executed by the model at each time step according to the order they are called. The corresponding description, the model entity executing the procedure, and the index of the compartmental transition (as illustrated in figure 2) are shown. The same procedure names are also used in the NetLogo program.

| Procedure | Description | Model entity | Compartmental transition |
|---|---|---|---|
| `update-the-current mosquito-capacity` | updates the variable that controls the seasonal density of adult mosquitoes. | nodes | - |
| `check-larval-density` | updates the daily survival parameter and the number of immatures in the breeding site. | nodes | - |
| `eliminate-adult-mosquitoes` | applies a mortality rate for each compartment of the mosquito adult population. | nodes | - |
| `from-newly-emerged to-blood-seeking` | adult mosquitoes in the newly-emerged compartment change to the blood-seeking compartment. | nodes | 1 - 2 |
| `mosquito-adult-emergence` | the newly-emerged compartment receives new adults emerging from the immature stage. | nodes | 3 - 4 |
| `from-gravid-to-blood-seeking` | gravid/pregnant mosquitoes lay their eggs and change from gravid to the blood-seeking compartment. | nodes | 5 - 8 |
| `from-blood-fed-to-gravid` | blood-fed mosquitoes that have matured their eggs change to the gravid compartment. | nodes | 9 - 11 |
| `from-exposed-to-infectious` | the exposed mosquitoes that reach the infectious stage change to the infectious compartment. | nodes | 12 - 16 |
| `vector-host-interaction` | blood-seeking mosquitoes that performed a successful bite on the hosts change to the blood-fed compartment. Susceptible blood-seeking mosquitoes and susceptible hosts that are infected after an interaction move to the blood-fed exposed compartment | nodes | 17 - 22 |
| `mosquito-dispersion` | the exchange between mosquitoes of linked nodes is performed, simulating the dispersal of mosquitoes. | nodes and links | 23 |

| | | | |
|---|---|---|---|
| `host-movement` | the exchange between hosts from linked nodes is performed, simulating host movement. | nodes and links | 24 |
| `host-recovery-or-death` | checks the infectious compartments of hosts and selects a number of them to be removed (die) or moved to the immune compartment. | nodes | 25 - 27 |
| `population-dynamics-of-hosts` | applies natural mortality and new births for the hosts. | nodes | - |
| `insert-the-virus` | at a given moment a proportion of susceptible mosquitoes become infectious in one or more external nodes to start the circulation of the virus in the landscape. | nodes | - |
| `calc-the- prob-of finding-a-host` | calculates the probability of blood-seeking mosquitoes finding a host in the next time step. | nodes | - |
| `calculate-speed` | calculates the spreading speed of the virus | observer | |
| `check-viral-circulation` | check if the virus is circulating in the mosquito or host populations. | nodes | - |
| `check stop conditions and update the time step` | the stop conditions are checked and the time step is updated. | observer | - |

#### 4. Design concepts

The basic concepts important to the design of the model are presented in this section. The model does not include the design concepts adaptation, objectives, learning and prediction.

##### 4.1. Basic Principles

The model addresses how spatial configurations of fragmented landscapes can influence the transmission and spread of vector-borne diseases (VBD). Particularly the model focus is on the dynamics of SYF in Brazil. By definition, VBDs are transmission systems involving at least three interacting species, a vector, a host, and a pathogen (Reisen, 2010). The loss and fragmentation of natural habitats can alter the transmission dynamics by causing changes in the abundance, behavior, and dispersal ability of the species involved in the transmission system (Estrada-Penã et al., 2014). In fragmented forest landscapes, the structure and connectivity of fragments can facilitate or hinder both the dispersion of organisms and interspecific contact rates, with direct consequences on pathogen transmission (McCallum and Dobson, 2002, Reisen, 2010). In this context, outbreaks of SYF in Brazil can be used as a study model to investigate the effect of landscape configuration on VBD spread.

Dispersal events of YFV from its endemic area in the Amazon to non-endemic extra-Amazonian regions of Brazil occur at least once every decade, causing epizootic and epidemic waves towards the East and South territories of the country (Franco, 1969;

Vasconcelos et al. 2001; Monath and Vasconcelos, 2015). From 2014 to 2021, Brazil experienced the largest SYF outbreak ever recorded in the country, with a high number of human cases and epizootics in non-human primates. The YFV has expanded from its endemic area to Brazilian Atlantic Coast, spreading through highly fragmented landscapes. The dispersion of YFV on the Atlantic Forest occurred through ecological corridors formed by remnants of forest fragments that serve as habitats for NHP (howler monkeys and marmosets) and vector mosquitoes (*Haemagogus* spp. and *Sabethes* spp.) (Possas et al., 2018; Brazilian Ministry of Health, 2019).

Recent studies based on data obtained from the epizootic-epidemic wave observed between 2017 – 2019 in forests of southeastern Brazil have suggested a possible relationship between highly fragmented areas with increased risk of disease in monkeys and humans and an easier permeability of YFV through the landscape (Ilacqua et al., 2021, Prist et al., 2022, Wilk-da-Silva et al., 2022). A possible explanation is that high border density of these fragmented landscapes favors an increased abundance of vector mosquitoes and reduced diversity of dead-end hosts, which increases the contact rates between vectors and monkeys and facilitates the virus amplification and propagation.

The transmission of the YFV occurs when an infectious mosquito (a female individual) feeds on the blood of a susceptible vertebrate host or, conversely, when a susceptible mosquito becomes infected by feeding on the blood of an infectious host. The blood that mosquitoes ingest from the host is needed to develop the eggs, which are deposited in aquatic larval habitats found by the females a few days after the blood meal. The eggs then hatch and release the larvae, which will develop in this aquatic environment and become new adult individuals in a few weeks. These newly emerged adults will allow the virus to continue circulating if they are already infected via vertical transmission (transmission from the gravid female to its progeny) or if they feed on the blood of infectious hosts (Medeiros-Sousa et al., 2022).

At the landscape level, the ways YFV uses to disperse over highly fragmented areas are not yet fully understood. The displacement of infected NHP (howler monkey) through their territories and forest corridors is undoubtedly a way of spreading the virus. However, the limited territory and traveling ranges and the inability to use extensive deforested areas to migrate from one fragment to another make NHP inefficient spreaders of the virus on highly fragmented landscapes (Hervé, 1986; Bicca-Marques and Freitas, 2010). On the other hand, infected mosquitoes can spread the virus over long distances. For example, it was demonstrated that *Haemagogus* mosquitoes have a high flight capacity, in some cases traveling distances over 11 km (Causey et al., 1950). In addition, there is a possibility that infected mosquitoes passively disperse between fragments being carried by the wind (Almeida et al., 2019).

##### 4.2. Emergence

The spread and persistence of the virus in the landscape, as well as the mortality of hosts, are unpredictable patterns that emerge from the different characteristics and behaviors assigned to the model entities and different landscape configurations. The reproductive dynamic of the vector is also an emerging process driven by different lower-level processes that include the interaction of mosquitoes with hosts to obtain a blood meal, the egg development time after blood ingestion, oviposition in breeding sites, larval development, and the emergence of new adult mosquitoes. In turn, the seasonal population abundance of mosquitoes and the movement of individuals between landscape fragments are patterns imposed by the model's rules to simplify the natural complexity inherent to modeling the processes responsible for these patterns. Likewise, the model does not include emergent behaviors of animal species involved, rather their development, demographic and infection rates are imposed but may depend on other biotic or abiotic variables.

##### 4.5. Sensing

At initialization, nodes detect the presence of other neighboring nodes, based on an assigned detection radius, and create links with them. Forest patches detect the minimum distance to a forest patch of a linked neighboring node and this information is used by the links to calculate the dispersal rate between two areas (see sections 5, 7.2.10, and 7.2.11). The dispersal rate is used by the model to define the flow of individuals between two linked forest areas in the landscape.

The breeding site senses the environmental conditions that determine whether newly adult mosquitoes can emerge (parameter *current-K*), thus controlling the total abundance of mosquitoes in the environment (see sections 7.2.1 and 7.2.5). This mechanism allows the reproduction of a pattern observed in nature where immatures take a longer time to develop during colder-drier periods when the mosquito abundance tends to be lower.

#### 4.6. Interaction

Model interactions are represented by the transitions described in Section 2.2 and by dispersal between neighboring nodes. The direct interactions between vector and host are the main mechanism of the model to simulate the transmission of the virus between individuals and obtaining blood meal by mosquitoes. In addition, mosquitoes interact with breeding sites by laying their eggs, and breeding sites interacts with the environment, allowing mosquito abundance control. A mediated interaction between mosquitoes is also represented and is based on intraspecific competition for resources, which occurs through a density-dependent control mechanism in breeding-site patches. Finally, nodes interact with each other via links that allow the exchange of individuals, representing the dispersal of mosquitoes and hosts through different forest areas in the landscape.

#### 4.7. Stochasticity

During model simulation, most transitions between compartments are based on calculated reference values that represent the expected number of individuals that will change from one status to another. These reference values are used as the mean ($\lambda$) of a random Poisson distribution to generate integers representing the effective number of individuals moving between two compartments. The submodels `vector-host-interaction`, `from gravid-to-blood-seeking`, `blood-fed-to-gravid`, `from-exposed-to-infectious`, `host-recovery-or-die`, and `population-dynamics-of-hosts` use Poisson distributions to represent the natural variability in these processes (see section 7.2).

To define which infectious main host will be removed, a random number between 0 and 1 is generated, if this number is below the mortality parameter for main hosts the individual is eliminated, if it number is above the individual is moved to the immune compartment. Finally, the choice of the first external node to initiate the virus circulation in the landscape also occurs randomly.

##### 4.8. Collectives

The only explicit collectives in the model are forest fragments and landscape units, formed by sets of patches to represent the spatial structure of the grid. These collectives directly affect the number and position of nodes and the creation of links between them, in addition to defining the abundance of vectors and hosts. Nodes are entities that represent communities of target species (hosts and vectors) that interact epidemiologically with other communities. Although the main hosts of sylvatic YF, howler monkeys, tend to live in groups (average 4 - 6 individuals), such collective social structures are not represented in the model.

##### 4.9. Observation

To visualize the patterns generated by the model, four different outputs are evaluated: 1) the YFV spreading speed (km/day), 2) the YFV persistence (time in days), 3) mortality of monkeys due to YFV infection, and 4) the maximum proportion of infected mosquitoes. The two first patterns are analyzed at a system level and the two last at a local level (for a single node). For model analyses, the mean and variance of 1) YFV spreading speed, 2) YFV persistence, 3) mortality of monkeys in the landscape due to YFV infection and 4) the proportion of nodes where the virus circulates are measured for different scenarios and sets of parameters.

The Graphical User Interface of the model allows the user to visualize the representation of the landscape and the nodes. Each forest area represented by the nodes assumes different colors depending on the virus circulation status: green if the virus has not emerged, red if the virus is circulating, and yellow after the end of the viral circulation.

#### 5. Initialization

In the model world, comprised of 121x121 patches, a landscape of 144 km² with forest fragments is generated. If real landscapes are simulated, the model uses vector layers of real landscapes with an extent of 12 x 12 km. The real landscapes simulated in the model were obtained from the Forestry Inventory of the State of São Paulo 2020 and is available at: https://www.infraestruturameioambiente.sp.gov.br/sifesp/inventario-florestal/. A second input shapefile contains a grid of 144 grid cells of 1 km² and is superimposed on the landscape, dividing it into several small landscape units where the model nodes are created. Nodes situated up to 10 patches (1 km) from the border of the grid are assigned as external nodes and will not be accounted for in model outputs. The total number of nodes created in the grid is variable and depends on 1) the landscape configuration (quantity, size, and shape of fragments), and 2) a function to merge small nodes into large ones. Nodes are merged if they have less than a given number of patches and since the nodes being merged belong to the same forest fragment. This function optimizes the number of nodes in the landscape and reduces the computational demand. During model initialization, we set the parameter `maximum-size-to-merge` to 20 (see Table 3) to define the maximum number of patches a node must have to be eligible to be merged with a node from a neighboring landscape unit.

To create links we set the parameter `maximum-distance-link` to values between 14 and 26 patches (1.4 – 2.6km) during model simulations, based on a presumed maximum daily flight distances for mosquitoes *Haemagogus* (see ODD- Additional Information text A1). Each created link calculates and stores as variables: 1) the distance between the nodes, 2) the minimum distance between the areas represented by the nodes, 3) the dispersion rate of mosquitoes between nodes (see section 7.2.10), and 4) the movement rate of hosts between nodes (see section 7.2.11).

The population of adult mosquitoes in a node is defined based on the assigned number of mosquitoes per hectare, and the total abundance is divided equally among the compartments for reproductive status (25% initialized in each of the four states). For immatures, we assume an initial abundance of 7.5 times the adult population (see ODD-Additional Information text A2, table A2 and figure A1), with the total number divided equally among the first twelve compartments of the immature list. The initial number of hosts in each node depends on the number of individuals per hectare assigned for each type of host. For main and alternative hosts these values were set to 0.5, while for dead-end hosts values between 0.7 and 1.3 were simulated. The parameter for the maximum proportion of host movement is set to 0.3 (Table 3).

The model assumes that the initial number of adult mosquitoes in the node is also the maximum seasonal carrying capacity and therefore the simulation starts on the day of the maximum abundance of mosquitoes. Values between 350 and 650 initial mosquitoes per hectare were simulated during model analysis. The parameter for maximum proportion of mosquito dispersal was set to values between 0.14 and 0.26 during simulations (Table 3).

All mosquitoes and hosts are initialized in the susceptible compartments. The initial day of virus introduction is assigned during initialization and is defined to simulate the YFV emergence and spread during a given season of the year. Four different days of virus introduction were simulated to representing the seasons: 182 (winter), 273 (spring), 365 (summer), 455 (fall). The initial burden of infected mosquitoes was varied between 0.5% to 6% of the population at the chosen nodes (Table 3, see more details in the section 7.14)

#### 6. Input data

The model does not use temporal input data, i.e., no time series of e.g. environmental drivers.

#### 7. Details

The model procedures listed in Table 2 are submodels performed sequentially during a time step. Their details and their empirical and theoretical backgrounds are described in this section. Some of the submodels were adapted from Medeiros-Sousa et al. (2022). A description of the parameters used in the model and their assigned values are presented in Table 3. The variables included in the equations are presented in Table 4, along with their descriptions and their corresponding names in the program code. Equation numbers are also used, as comments, in the corresponding NetLogo code. Please note that all transitions described in the following are referring to mosquitos and hosts in a given node or, in the case of dispersal, between two given nodes.

**Table 3**. Model parameters and their values or range of values assigned during the model simulations. The description and assignment sources are also presented.

| Input parameters | Description | Value/range | Source |
|---|---|---|---|
| temporal resolution | the time represented by each time step | 1 day | assumed |
| spatial resolution | the area represented by each patch in the grid | 1 hectare (100m x 100m) | assumed |
| `k-max` | the maximum seasonal value for the abundance of mosquitoes (the maximum carrying capacity - current-K) in a node | initial-mosquitoes-per-ha * grid size of the node | sensitivity analysis |
| `k-min` | the minimum seasonal value for the abundance of mosquitoes (the minimum current-K) in a node | `k-max` / 6 | Chadee et al., 1992 |
| `alpha` | base coefficient in the equation of current-K | (`k-max` + `k-min`) / 2 | fitted |
| `beta` | trend coefficient in the equation of current-K | - (`k-max` - `k-min`) / (365 / 2) | fitted |

| | | | |
|---|---|---|---|
| gamma | seasonal coefficient in the equation of current-K | α – k-max | fitted |
| lv1 | maximum number of larvae in the breeding site in which the survival probability is >= 99% | 1 * k-max | fitted |
| lv2 | number of larvae in the breeding site in which the survival probability is 50% | 7.5 * k-max | fitted |
| time to start blood-seeking | time required for newly emerged mosquitoes to move to blood-seeking status (time steps) | 1 | Forattini, 2002 |
| time-to-egg-maturation | time required for engorged mosquitoes to maturate the eggs and start looking for breeding sites to lay eggs (time steps) | 5 | Dégallier et al., 1998 |
| extrinsic-incubation-period | time of extrinsic incubation period in an infected mosquito (time steps) | 12 | Staples and Monath, 2008; Johansson et al., 2010 |
| main-host-per-ha | the initial population of main hosts (Monkeys) per patch (assuming each patch of the grid represents one ha) | 0.5 | Bica-Marques et al., 2018 |
| latent-period | time elapsed from the infection to infectiousness in the vertebrate hosts (time steps) | 3 | Moreno et al., 2015 |
| main-host-viremic-period | the period when an infected main host is infectious to susceptible mosquitoes (time steps) | 5 | Moreno et al., 2015 |
| infectious-main-host-mortality | the probability a main host has of dying when infected with YFV (assuming the main hosts are howler monkeys) | 0.9 | Possamai et al., 2022 |
| biting-success | The probability of a blood-seeking mosquito having success (biting the host) during an interaction with a vertebrate host | 0.5 | assumed |
| daily-survival-rate | the probability that a given individual of the mosquito population will be alive at the 17ndo f a day | 0.93 | Estimated based on Marcondes and Alencar, 2010; Hervé et al., 1985 |
| mortality-rate | the probability that a given individual of the mosquito population will die during a time step | 1 - daily-survival-rate | Estimated based on Marcondes and Alencar, 2010; Hervé et al., 1985 |
| minimum-development-time | the minimum time required for new adult mosquitoes to emerge after the oviposition event (time steps) | 12 | Alencar et al., 2008 |
| vertical-infection-rate | the probability of a newly-emerged adult from a progeny of an infectious mosquito being infectious | 0.01 | Aitken et al., 1979; Dutary & Leduc, 1981; Lequime and Lambrechts, 2014; Alencar et al., 2021 |

| | | | |
|---|---|---|---|
| initial-mosquitoes-per-ha | the initial number of mosquitoes per forest patch (assuming each patch of the grid represents one ha) | –350 - 650 | sensitivity analysis |
| dead-end-host-per-ha | the population size of dead-end-hosts per hectare | 0.35 – 0.65 | sensitivity analysis |
| mosquito-trans-competence | the probability of an infectious mosquito will transmit the virus to a susceptible vertebrate host during an interaction | 0.2 | estimated based on Couto-Lima et al., 2017 |
| main-host-trans-competence | the probability of an infectious monkey will transmit the virus to a susceptible mosquito during an interaction | 0.6 | estimated based on Couto-Lima et al., 2017, Mares-Guia et al., 2020, and Fernandes et al, 2021 |
| prob-find-breed | the probability that a gravid mosquito will find a breeding site to lay its eggs within a time step | 1 | assumed |
| eggs-by-gravid-mosquito | the number of eggs a gravid/pregnant female of the vector species (*Haemagogus* ssp. in this case) will laid in the breeding site, considering only those eggs that will become adult females | 15 | Chadee and Tikasingh, 1989 and Mondet 1997 |
| YFV-initial-burden | the proportion of mosquitoes that will change their status from susceptible to infectious in the chosen external nodes to initiate the transmission dynamics in the landscape | 0.005 - 0.04 | sensitivity analysis |
| max-prop-mosq-dispersal | the maximum proportion of mosquitoes allowed to disperse to a linked node during a time step | 0.14 - 0.26 | sensitivity analysis |
| host-move-proportion | the maximum proportion of hosts allowed to move to a neighbor node during a time step | 0.3 | sensitivity analysis |
| alternative-host-per-ha | the initial population of alternative hosts per patch (assuming each patch of the grid represents one ha) | 0.5 | assumed |
| main-host-mean-lifespan | the average time of life in days for a howler monkey (time steps) | 5475 | Buss et al., 2019 |
| alt-host-mean-lifespan | the average time of life in days for an alternative host (time steps) | 3650 | assumed |
| alt-host-viremic-period | the period an infected individual is infectious to a susceptible mosquito (days) | 5 | assumed |
| alt-host-trans-competence | the probability of an infectious alternative host transmitting the virus to a susceptible mosquito during an interaction | 0.2 | assumed |
| external-area | the area in patches from the border of the grid that is defined as external to the landscape | 10 | assumed |
| maximum-size-to-merge | the maximum size in patches for one node to be eligible to merge with another | 20 | assumed |
| maximum-distance-link | the maximum distance in patches between two nodes to create a link | 14 - 26 | sensitivity analysis |

| | | | |
|---|---|---|---|
| `start-YFV-circulation` | The day after the initialization the virus will start circulating in the landscape | 183 – 455 | sensitivity analysis |

**Table 4.** Variables present in the equations of the submodels, their descriptions, and corresponding names in the program code.

| Variable | Description | Assigned name in the program code |
|---|---|---|
| *current-K* | seasonal carrying capacity of mosquito population | `current-k` |
| $P$ | the daily survival probability of immatures | `P` |
| $im_{tot}$ | the total number of immatures in the breeding site | `tot-immat` |
| $n_S^{ne}$ | susceptible individuals added to the newly-emerged compartment | `new-emerged-susc` |
| $n_I^{ne}$ | infectious individuals added to the newly-emerged compartment | `new-emerged-infe` |
| $p_{im}$ | the number of potentially infectious newly-emerged mosquitoes | `num-emerged-infec-mosq` |
| $n_i^{bs}$ | number of blood-seeking mosquitoes in the epidemic compartment *i* (susceptible, exposed, or infectious) | `susc-bs`<br>`expo-bs`<br>`infe-bs` |
| $n_i^{bf}$ | number of blood-fed mosquitoes in the epidemic compartment *i* | `susc-bf`<br>`expo-bf`<br>`infe-bf` |
| $n_i^{gr}$ | number of gravid mosquitoes in the epidemic compartment *i* (susceptible, exposed, or infectious) | `susc-gr`<br>`expo-gr`<br>`infe-gr` |
| $n_E^i$ | number of exposed individuals in the reproductive compartment *i* | `expo-bs`<br>`expo-bf`<br>`expo-gr` |
| σ | the interaction probability between a blood-seeking mosquito and a host | `interactions` |
| $p_h$ | the probability that a blood-seeking mosquito will find a host within a time step | `prob-find-host` |
| $M_{ne}$ | the daily referential number of newly exposed/infected mosquitoes | `new-infected-mosq` |

| | | |
|---|---|---|
| $P_{ne}$ | the daily referential number of newly exposed/infected main hosts | `new-infected-main-host` |
| $A_{ne}$ | the daily referential number of newly exposed/infected alternative hosts | `new-infected-alt-host` |
| $H$ | total number of hosts | `tot-host` |
| $P$ | number of main hosts | `current-main-host-num` |
| $A$ | number of alternative hosts | `current-alt-host-num` |
| $P_S$ | number of susceptible main hosts | `susc-main-host` |
| $A_S$ | number of susceptible alternative hosts | `susc-alt-host` |
| $B_h$ | daily number of bite per host | `num-bites-per-host` |
| $M_{disp}$ | the dispersal rate of mosquitoes or host movement rate between two linked nodes | `dispersal-rate-end1` `dispersal-rate-end2` `host-movement-rate-end1` `host-movement-rate-end2` |
| $nd_{sz}^{1}$ | the size in patches of node end1 | `size-end1` |
| $nd_{sz}^{2}$ | the size in patches of node end2 | `size-end2` |
| $n_I^{mh}$ | number of infectious main hosts | `infe-main-host` |
| $n_I^{ah}$ | number of infectious alternative hosts | `infe-alt-host` |
| $n_R^{mh}$ | number of recovered or immune main hosts | `infe-mh-to-rec-death` |
| $dead$ | number of main hosts that die and are removed | `dead-mh` |

#### 7.1. Update the current mosquito capacity

The abundance of mosquitoes, including those of the genus *Haemagogus*, is influenced by seasonal variations in rainfall, temperature, and relative humidity (Marcondes and Alencar, 2010; Ribeiro et al., 2012; Couto-Lima et al., 2020). For example, Chadee et al. (1992) observed that the abundance of *H. janthinomys* in Trinidad was up to six times greater in rainy periods than in dry periods. To simulate the effect of seasonal variations on mosquito populations, each node has a state variable that controls the local seasonal carrying capacity of mosquitoes, here called 'current-*K*' (Figure 3). Current-*K* has two purposes: (1) to simulate the effect of seasonal variations in the abundance of adult mosquitoes during a year (i.e., 365 time steps in the temporal resolution of the model); (2) to control the emergence of new adult mosquitoes as the transition of individuals from the immature list to the adult list occur only when the adult abundance is below the *current-K*.

*Current-K* is updated at each time step with the following equation:

$$\textit{current-K} = \alpha + \beta \sin\left(2\pi t\left(\frac{180}{\pi}\right)\right) + \gamma \cos\left(2\pi t\left(\frac{180}{\pi}\right)\right), \quad \text{(Eq. 1)}$$

where:

$\alpha$ = (`k-max` + `k-min`) / 2

`k-max` = initial number of mosquitoes in the node

`k-min` = `k-max` / 6

$\beta$ = – (`k-max` – `k-min`) / (365 / 2)

γ = $\alpha$ – `k-max`

$t$ = 0.5 + (time / 365).

The sine and cosine terms assign a cyclic pattern to the equation (Figure 3). The constant 2π(180/π) is the circumference of a circle, where 180/π does the conversion from degrees (NetLogo standard) to radians.

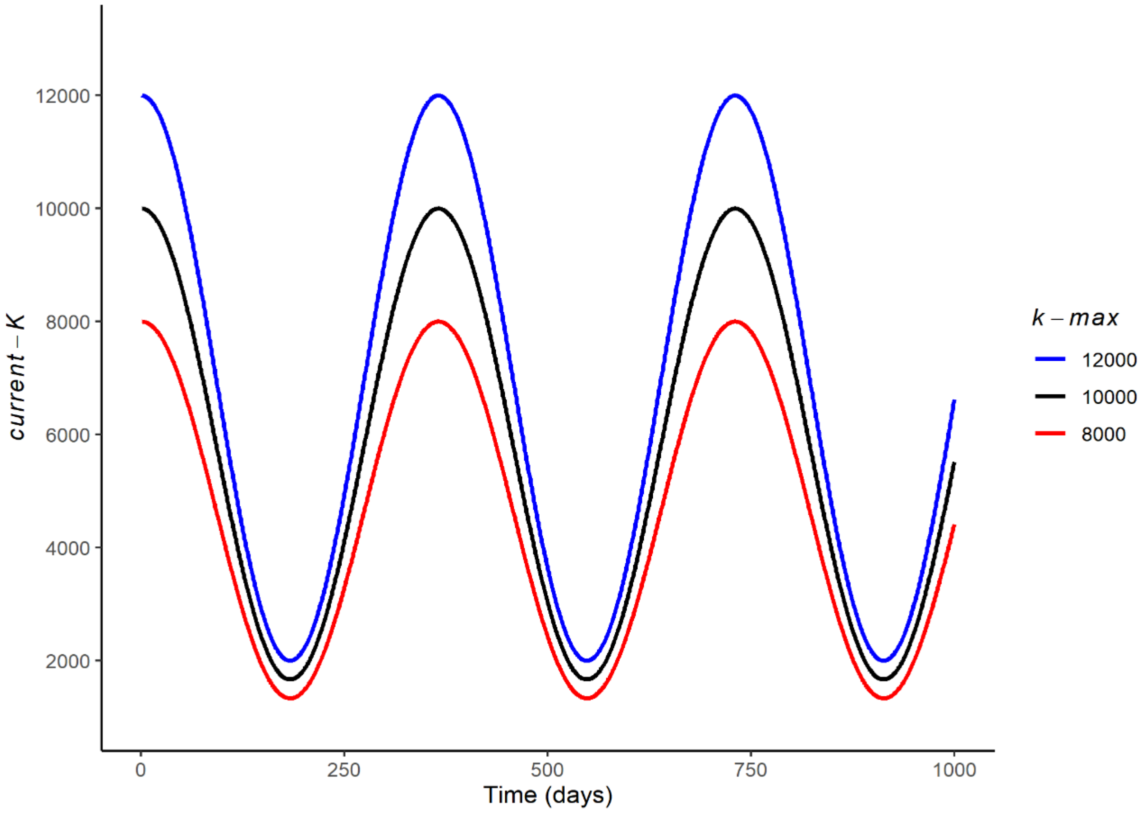


**Figure 3.** Illustration of the cyclic pattern of *current-K* as a function of time and for three different values of *k-max*.

#### 7.2. Check larval density

Mosquitoes *Haemagogus* usually lay their eggs in tree holes and bamboo internodes (Marcondes and Alencar, 2010). The model represents these breeding places as a single breeding site. Each node has its breeding site capacity based on two lists containing information about the immatures and their development stage. The first list contains the total number of immatures in the area and the order of the compartments represent the time of development of these individuals, as illustrated in Figure 2B. The second list represents the number of possible infected immatures from each compartment of the total immatures list (see sections 7.5 and 7.6). Therefore, each compartment on the first list has a corresponding compartment in the second list.

At each time step, the node checks the immature lists and eliminates a proportion of individuals from each compartment based on a rate *P* (daily survival probability of immatures), which decreases as the larval density increases. This mechanism aims to simulate density-dependent intraspecific competition (Walker et al., 1991). The variable *P* follows a logistic distribution (Figure 4) and is obtained from four parameters representing two arbitrary points in a logistic curve: $sv_1$ = 0.99, $sv_2$ = 0.5, `lv1` = 1 * *k-max*, and `lv2` = 7.5 * *k-max*. The $sv_1$ represents the daily survival probability (value of y) when the abundance of immatures is equal to `lv1` (value of x) and *sv2* represents the daily survival probability when the abundance of immatures is equal to `lv2`. We opted for values $sv_1$ = 0.99 and $sv_2$ = 0.5 to represent a point when the survival is very high (99%) and a second point when the survival and death probabilities are equal (50%) (Figure 4). As higher the values assigned to `lv1` and `lv2` more individuals will survive and become adults. As more adults emerge, the carrying capacity is reached more frequently, leading to more retention of immature individuals in the breeding site and increasing the time they will take to become adults. Therefore, `lv1` and `lv2` have influence on the average minimum and maximum larval development times. Thus, the parameters `lv1` and `lv2` were calibrated to reproduce plausible patterns based on empirical values for immature development time so that this time should be short in favorable seasonal periods and relatively long in unfavorable seasonal periods (Alencar et al., 2008; Marcondes and Alencar, 2010) (see ODD- Additional Information text A2 and figure A1). Based on these parameters, the immatures daily survival rate ($P$) is calculated as follows:

$D = ln\ (sv_1 / (1- sv_1))$ (Eq 2.1)

$C = ln\ (sv_2 / (1- sv_2))$ (Eq 2.2)

$B = (D – C) / (lv_1 - lv_2)$ (Eq 2.3)

$A = D – B * lv_1$ (Eq 2.4)

$Z = exp\ (A + B * im_{tot})$ (Eq 2.5)

$P = Z / (1+Z)$. (Eq 2.6)

Where $im_{tot}$ represents the current total number of immatures in the breeding site.

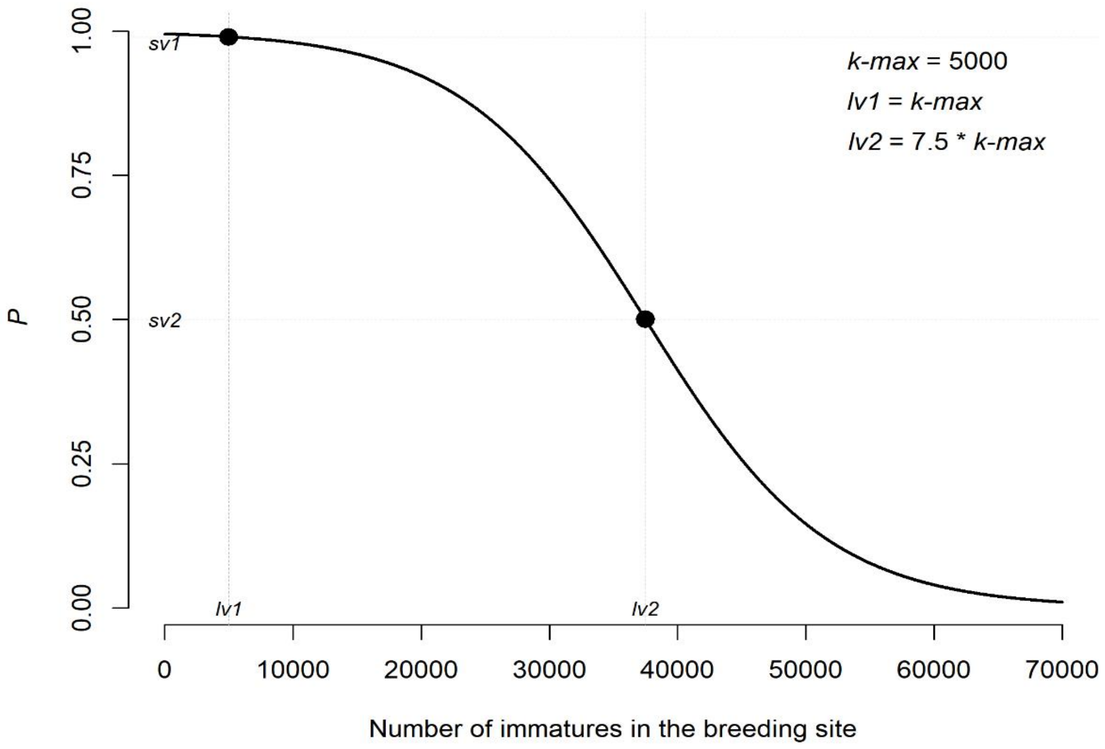


**Figure 4**. Daily survival probability of immatures (*P*) as a function of the number of immatures in the breeding site. Black dots indicate the value of $lv_1$ and $lv_2$ in x and $sv_1$ and $sv_2$ in y.

##### 7.3. Eliminate adult mosquitoes

A mortality function is applied for each list of adult mosquitoes in the nodes. Each compartment of the list has its number of individuals multiplied by a constant daily (finite) mortality rate (μ). From the obtained number, 0.5 is subtracted and the result is rounded to the closest integer. The subtraction by the constant 0.5 ensure the compartment number can go to 0 when $\mu > 0.5$. The result indicates the number of individuals to be eliminated from the compartment at the time step. The value of μ is calculated based on the `daily-survival-rate` (S) for *Haemagogus* using the equation $\mu = 1 - S$ (Eq. 3). Studies with *Haemagogus* mosquitoes have estimated a daily survival rate of between 0.9 and 0.96 (Marcondes and Alencar, 2010). Here we assumed S = 0.93, following the calibration analysis performed by Medeiros-Sousa et al. (2022).

##### 7.4. *From newly emerged to blood-seeking* (compartmental transitions 1 and 2)

In each time step all adult mosquitoes in the newly-emerged compartments change to respective blood-seeking compartments. This waiting period of one day for newly-emerged mosquitoes is included in the simulation to take into account that after the emergence the mosquito needs about 24 hours for the exoskeleton to harden and for individuals to mate (Forattini, 2002).

##### 7.5. *Mosquito adult emergence* (compartmental transitions 3 and 4)

The total number of adult mosquitoes in the node is accounted for. If the adult abundance is below the carrying capacity the emergence of newly adult mosquitoes is allowed to occur. Immatures are transferred to adult newly emerged compartment, starting by older immature compartments, if the number of adults is below the carrying capacity. All the individuals in the compartment are moved, even if only part fits into the capacity. Only individuals with a time equal to or greater than the `minimum-development-time`, assumed to be 12 days (Alencar et al., 2008), are allowed to become adults. Thus, once the immature list has been reduced to individuals with less than 12 days the emergence stops, even if the carrying capacity has not been reached.

Since infectious females of *Haemagogus* mosquitoes can transmit the virus to their progeny via vertical/transovarial transmission (Mondet et al., 2002), we include a mechanism where each compartment in the immature list has a corresponding compartment in a second list that accounts for the number of potentially infected immatures (see section 7.6). For each compartment removed from the immature list and added to the newly emerged compartment ($n^{ne}$), a correspondent compartment is removed from the potentially infected list and accounted as a possible infectious newly emerged. From the total newly emerged in the time step certain proportions will be susceptible $n_S^{ne}$ or be infectious $n_I^{ne}$. To obtain $n_I^{ne}$,, a random-Poisson function with $\lambda = p_{im}VIR$ (Eq. 4) is used, where $p_{im}$ is the number of potentially infectious newly emerged mosquitoes and $VIR$ is the parameter `vertical-infection-rate`. The generated random value represents $n_I^{ne}$ and the $n_S^{ne} = n^{ne} - n_I^{ne}$.

The frequency of vertical transmission observed exprimentally for most arboviruses, including the YFV, is generally low (< 1%) (Aitken et al., 1979; Dutary & Leduc, 1981; Lequime and Lambrechts, 2014). Therefore, we assumed a VIR = 0.01.

##### 7.6. *From gravid to blood-seeking* (compartmental transitions 5, 6, 7, and 8)

The number of individuals moving from gravid to the blood-seeking compartment is obtained by a random-Poisson distribution with $\lambda = n_i^{gr} p_{fb}$ (Eq. 5), where $n_i^{gr}$ is the number of gravid mosquitoes in compartment $i$ for epidemic status (SEI) and $p_{fb}$ is the probability of a gravid mosquito finding a breeding site to lay eggs (parameter `prob-find-breed`). The generated random value represents the gravid females that laid their eggs and moved to the blood-seeking compartment. For each gravid mosquito that moves to blood-seeking, the model adds a number of individuals (representing the eggs) to the first compartment of the immature list. The eggs laid by gravid mosquitoes in the infectious state will be added to both the first compartment of the immature list and the first compartment of the potentially infected immature list.

Oviposition is interpreted here as the deposition of the entire egg load of a gravid female in a gonotrophic cycle, which ranges from 15 to 40 eggs for *Haemagogus* (Chadee and Tikasingh, 1989, Mondet 1997). For model purposes we assumed that a gravid female lays 30 eggs in her gonotrophic cycle, half of which will become adult females. Since only females are represented in the model, each gravid mosquito adds 15 eggs per gonotrophic cycle in the immature list. For simplicity, we consider well-distributed and abundant breeding sites in the forest fragments and therefore we assigned $p_{fb}$ = 1.

##### 7.7. *From blood fed to gravid* (compartmental transitions 9, 10, and 11)

A proportion of mosquitoes in the blood-fed status is moved to the gravid compartment based on the parameter `time-to-egg-maturation`. The number of individuals that will move is obtained by a random-Poisson distribution with $\lambda = n_i^{bf} \frac{1}{mt}$ (Eq. 6), where $n_i^{bf}$ is the number of blood-fed individuals in the epidemic compartment $i$ and $mt$ is `time-to-egg-maturation`. The time to egg maturation after the blood meal was assumed to be 5 days, as for *Haemagogus* gonotrophic cycles from 5 to 7 days have been observed (Mondet 1997; Dégallier et al., 1998).

##### 7.8. *From exposed to infectious* (compartmental transitions 12, 13, 14, 15, and 16)

After being infected, mosquitoes and hosts have a period in which the virus starts replicating in the body but is still at very low levels to be transmitted to other individuals. During this period, the individual is considered exposed and infected by the virus but is not yet infectious to other individuals. For mosquitoes, this period is called extrinsic incubation and comprises the time taken for the virus to replicate and disseminate to the salivary glands, while for vertebrate hosts it is the intrinsic incubation or latent period, preceding the infectiousness and first symptoms (Johansson et al., 2010).

To simulate the transition from the exposed to infectious state we use a random-Poisson distribution with $\lambda = n_E^i \frac{1}{ep}$ (Eq. 7) for mosquitoes, and $\lambda = n_E^i \frac{1}{lp}$ (Eq. 8) for hosts, where, $n_E^i$ is the number of exposed individuals in compartment *i*, *ep* is the `extrinsic-incubation-period`, and *lp* is the `latent-period`. The value obtained is subtracted from the compartment of exposed individuals and added to a respective compartment of infectious. Based on experimental studies with the vector *Aedes aegypti* (Johansson et al., 2010), the extrinsic incubation period for the YFV was assumed to be twelve days. In turn, a latent period of 3 days is assumed for both monkeys and alternative hosts (Moreno et al., 2015).

#### 7.9. *Vector-host interaction* (compartmental transitions 17, 18, 19, 20, 21, and 22)

Mosquitoes feed primarily on plant nectar, but females also need to ingest blood to ensure the proteins and nutrients necessary for the development of their eggs. This blood-feeding behavior allows the transmission and maintenance of pathogens between mosquitoes and vertebrate animals when an infectious mosquito feeds on the blood of a susceptible vertebrate host or, conversely, when a susceptible mosquito becomes infected by feeding on the blood of an infectious host (Clements, 1993; Forattini, 2002).

To simulate this interaction between mosquitoes and hosts in the model, we calculate four values: 1) the mosquito interaction probability with hosts (σ), 2) the daily referential number of newly exposed/infected mosquitoes ($M_{nE}$), 3) the daily referential number of newly exposed/infected main hosts ($P_{nE}$), and 4) the daily referential number of newly exposed/infected alternative hosts ($A_{nE}$). The referential numbers are used as parameters for random-poisson distribution to generate stochastic values.

Only blood-seeking mosquitoes will interact with hosts during a model time step, and the probability they interact is calculated as:

$$\sigma = p_h b \quad \text{(Eq. 9)}$$

Where $b$ is the daily `biting-success`, assumed to be 0.5, and $p_h$ is the probability of a blood-seeking mosquito finding a host (see section 7.15). Hence, the number of individuals from a blood-seeking compartment $i$ ($n_i^{bs}$) moving to blood-fed compartment $i$, ($n_i^{bf}$), is obtained using a random-Poisson distribution with $\lambda = n_i^{bs}\sigma$ (Eq. 10).

Among the susceptible mosquitoes moving to the blood-fed compartment, some can be infected when they 'bite' an infectious host. Considering the interaction with main hosts, for example, the daily referential number of newly exposed mosquitoes ($M_{nE}$) will depend on: 1) the number of susceptible blood-seeking mosquitoes ($n_S^{bs}$), 2) the probability of these mosquitoes interacting with a host in the time step ($\sigma$), 3) the proportion of main hosts among the total number of hosts ($\frac{P}{H}$), 4) the proportion of infectious individuals among the main hosts ($P_I$), and 5) the transmission competence of the infectious main hosts ($\delta_P$; parameter `main-host-trans-competence in Table 3`). The same logic can be applied when we consider the interaction of susceptible mosquitoes with alternative hosts (A). Thus, the referential daily number of newly exposed blood-fed mosquitoes is calculated as:

$$M_{nE} = n_S^{bs}\sigma\frac{P}{H}P_I\delta_P + n_S^{bs}\sigma\frac{A}{H}A_I\delta_A \quad \text{(Eq. 11)}$$

The term $n_S^{bs}\sigma$ is obtained from a random-Poisson distribution, as explained above.

For main hosts, the referential daily number of newly exposed individuals ($P_{nE}$) will depend on: 1) the number of susceptible main hosts ($P_S$), 2) the daily number of bites per host ($B_h$), 3) the infectious blood-seeking over the total blood-seeking ($\frac{N_I^{bs}}{N^{bs}}$), and 4) the transmission competence of infectious mosquitoes ($\delta_m$; parameter `mosquito-trans-competence` in Table 3). The referential daily number of newly exposed main hosts is calculated as:

$$P_{nE} = P_S B_h \frac{N_I^{bs}}{N^{bs}}\delta_m \quad \text{(Eq. 12)}$$

Where, $B_h = \frac{\sigma N^{bs}}{H}$ (Eq. 13). For alternative hosts ($A_{nE}$) the variable $P_S$ is changed by $A_S$.

After obtaining these reference daily numbers for mosquitoes and hosts, these values are used to generate the "real" (stochastic) daily numbers of exposed individuals, using a random-Poisson distribution ($\lambda = M_{nE}$; $\lambda = P_{nE}$; $\lambda = A_{nE}$). Finally, the values are added to their respective new compartments and subtracted from their old compartments. We assume $\delta = 0.20$ for mosquitoes, $\delta = 0.6$ for main hosts and $\delta = 0.2$ for alternative hosts

(Couto-Lima et al., 2017; Mares-Guia et al., 2020; see Table 3 and ODD- Additional Information text A3).

##### 7.10. *Mosquito dispersion* (compartmental transition 23)

Mosquitoes are considered the main dispersers of the YFV in the forests (Possas et al., 2018; Abreu et al., 2022). *Haemagogus janthinomys*, for example, were collected at distances up to 11.5 km from a releasing point in southeast Brazil (Causey et al., 1950). Here we assume that the dispersal of mosquitoes is a function of the minimum distance between two linked node areas and the proportional size of them in relation to a landscape unit. The dispersal rate of mosquitoes ($M_{disp}$) between two linked nodes is calculated by the link during the model initialization using the following equations:

$M_{disp}^{1\to2} = \frac{1}{\min dist} max_{disp} \frac{nd_{sz}^{2}}{lg_{sz}}$ (Eq. 14) and $M_{disp}^{2\to1} = \frac{1}{\min dist} max_{disp} \frac{nd_{sz}^{1}}{lg_{sz}}$ (Eq. 15)

Where, $M_{disp}^{1\to2}$ is the dispersal rate from node 1 to node 2, $M_{disp}^{2\to1}$ is the dispersal rate from node 2 to node 1, $\min dist$ is the minimum distance between the two node areas, $max_{disp}$ is the parameter for the maximum dispersal of mosquitoes between two nodes, $nd_{sz}^{1}$ and $nd_{sz}^{2}$ are the size in patches of the forest fragements represented by the two linked nodes (i.e., the node variable `my-grid-size`), and $lg_{sz}$ is the size in patches of the landscape unit. The $\frac{1}{\min dist}$ promotes a linear reduction of mosquito dispersal between nodes as $\min dist$ increases. The term $\frac{nd_{sz}}{lg_{sz}}$ allows the dispersal to be proportional to the relative size of the node. In the current configuration of the model the $lg_{sz} = 100$ patches. The parameter $max_{disp}$ (`max-prop-mosq-dispersal` in Table 3) represents the maximum proportion of mosquitoes that disperse to the linked node during a time step, which occurs when $\min dist = 1$ and $\frac{nd_{sz}}{lg_{sz}} = 1$.

To define the mosquitoes that will move between linked nodes the number of individuals in each compartment of nodes 1 and 2 are multiplied by $M_{disp}^{1\to2}$ or $M_{disp}^{1\to2}$, respectively. After this, each node subtracts the number of dispersed individuals from their respective compartments and adds the number of migrants coming from the linked node. Only individuals in the newly emerged compartment are assumed not to disperse since these individuals need to harden the exoskeleton before being able to fly.

If an exposed or infectious individual arrives at the node, a new virus emergence event is reported.

##### 7.11. *Host movement* (compartmental transition 24)

The displacement of howler monkeys through their territories and forest corridors is one of the ways of spreading the virus in the landscape. However, these animals have small territory and tend to not move through extensive deforested areas to migrate from one fragment to another (Hervé, 1986; Bicca-Marques and Freitas, 2010). Based on these observations, the model assumes the movement of main hosts in the landscape is limited to linked nodes with minimum distance equal or lower than 2 patches (200 m). For simplicity, the same dispersion limitation was assumed for alternative and dead-end hosts. The entire procedure is performed using the same sequence described for mosquito dispersion, with two main differences: 1) the links will change $max_{disp}$ by the parameter `host-move-proportion` to calculate $M_{disp}^{1 \to 2}$ and $M_{disp}^{2 \to 1}$ (now defined as 'host movement rate between nodes'), 2) the function to exchange individuals between linked nodes is applied for all host compartments, except for infectious main hosts since howler monkeys tend to stop moving in the forest after the disease onset.

##### 7.12. *Host recovery or death* (compartmental transitions 25, 26, and 27)

Infectious hosts are those that present enough viremia in the bloodstream to infect susceptible mosquitoes that bite them. The viremic period for YFV will cease after a few days either by the death of the individual or by a response of the immune system that leads to recovery and immunity to the virus (Engelmann et al., 2014, Fernandes et al., 2021).

A random-Poisson distribution is used to obtain the number of individuals that will move from the infectious compartment and become immune or be removed from the simulation. The lambda parameter is calculated as: $\lambda = n_I^{mh} \frac{1}{v_{mh}}$ (Eq. 17) for main hosts and $\lambda = n_I^{ah} \frac{1}{v_{ah}}$ (Eq. 18) for alternative hosts. Where, $n_I^{mh}$ and $n_I^{ah}$ are the number of infectious main hosts and alternative hosts, and $v_{mh}$ and $v_{ah}$ are the parameters `main-host-viremic-period` and `alt-host-viremic-period`, respectively.

For alternative hosts, the obtained value by the random-Poisson distribution represents the recovered and immune individuals since mortality is disregarded for these hosts in the model. Thus the value is directly subtracted from the infectious compartment and added to the immune compartment, in turn, the main hosts can recover from the disease and become immune or can die and be removed from the model. Therefore, from the random value obtained for recovered/removed ($n_R^{mh}$) main hosts the number of dead or surviving individuals is defined using the following steps: 1) a list with length = $n_R^{mh}$is created and for each element is assigned a random real number between 0 and 1 generated from a uniform distribution, 2) a function is applied to report the number of elements lower than the `infectious-main-host-mortality`, the result representing the *dead* individuals, 3) the $n_R^{mh}$ is subtracted from the infectious compartment and $n_R^{mh} - dead$ is added to the immune compartment.

The viremic period assumed for howler monkeys is five days (Rodhain, 1991; Bicca-Marques and Freitas, 2010). For simplicity, we considered the same viremic period for alternative hosts. The probability of death for infectious howler monkeys was set to 0.9 as high mortality tends to be observed for these animals during epizootic events of YFV (see Possamai et al., 2022).

#### 7.13. Population dynamics of hosts

The mortality of howler monkeys and alternative hosts by causes not linked to YF as well as the births are considered in the model. In the current version, a simplified model of population dynamics is implemented and for each individual that dies in a given node a new susceptible individual is born in the same node, keeping the population size constant in the landscape in the absence of the virus. The number of individuals that will die in the time step is obtained by a random-Poisson distribution with lambda calculated as: $\lambda = n_i \frac{1}{lp}$ (Eq. 19). Where, $n_i$ is the number of individuals in compartment $i$ and $lp$ is the expected lifespan for a individual of the population (`main-host-mean-lifespan` and `alt-host-mean-lifespan` in Table 3). The number of individuals that die is subtracted from its respective compartment and added to the susceptible compartment to simulate new births.

For howler monkeys, we assume $lp =$ 5475 days (15 years) following field observations (Buss et al., 2019). For alternative hosts, variable values for $lp$ parameter can be assigned depending on the populational turnover assumed in the simulation. As default, the model assumes a lifespan of ten years (3650 days) for alternative hosts.

Since one of the goals of the study is to verify the impact of YFV on the initial population of howler monkeys in the landscape, dynamics of population replacement and recolonization of fragments by these animals after epizootic events were not included in the model.

#### 7.14. Insert the virus

At a certain point in the simulation, the virus is introduced in the landscape via infectious mosquitoes. This procedure can be compared to an imaginary experiment in which many infectious mosquitoes are released into some initial points and outcomes of interest are observed in the landscape. Two parameters control the virus insertion in the landscape: 1) the `start-YFV-circulation` that indicates the day the first infectious individuals will be introduced in the landscape, and 2) the `YFV-initial-burden`, that defines the proportion of infectious mosquitoes introduced per selected node (see below).

First, it is checked if the simulation is on the day of virus introduction defined by `start-YFV-circulation`, if so, a first external nodes is selected randomly and its status for the boolean variable `initiator` is changed to `true`. This first initiator node selects other four neighbor external nodes that also change their status for `initiator = true`, forming a spatial cluster of neighbor initiator nodes. For each initiator node, the number of susceptible adult mosquitoes in each reproductive compartment is multiplied by the `YFV-initial-burden`. The result is a new list containing the new infectious individuals. The values of this new list are subtracted from the susceptible compartment and included in the infectious compartment. The ‘release’ of infectious mosquitoes in five nodes reduces the bias of initial node choices by selecting nodes representing forest areas of different sizes and abundance of mosquitoes. Furthermore, the virus introduction simulates a 'wave-front' of the virus that arrives in an area in the limits of the landcape (external nodes).

#### 7.15. Calculating the probability of finding a host

The procedure uses a fitted Michaelis-Menten function to calculate and update the probability of a mosquito finding a host during the time step, as follows:

$$p_h = 2.382 * \frac{Total\ hosts}{107.125 + Total\ hosts} \quad \text{(Eq. 20)}$$

For model purposes $p_h > 1 = 1$. The parameter values of the function were fitted using outputs of the model proposed by Medeiros-Sousa et al. (2022) (See ODD- Additional Information text A4 and Figure A2).

#### 7.16. Calculate speed

This procedure is responsible for calculating the average daily distance traveled by the virus within a given time interval. For this, a point called centroid-initiator is obtained, which is the centroid of the cluster of patches belonging to the initiator nodes and represents the point where the virus began to circulate in the landscape. The distance from this point to the most distant node to which the virus has spread is then calculated and this distance is divided by the number of days elapsed since the emergence of the virus, thus obtaining the speed of propagation.

#### 7.17 Check viral circulation

In each time step all the nodes update their number of infected individuals (sum of infected adult mosquitoes and infected hosts). Nodes with active transmission are asked to check both the number of infected individuals and the number of potentially infected immatures. If both present zero individuals then the node changes the active transmission status to false.

#### 7.18. Check stop conditions and update the time step

Three criteria are needed to stop the simulation: (1) the time step must be greater than the parameter `start-YFV-circulation`; (2) there must be no mosquitoes or hosts in the infectious or exposed state in the landscape; and (3) there must be no oviposition events by infectious mosquitoes in the landscape. Once these three conditions are met, the simulation is completed since the virus is no longer circulating in the environment. If the stop conditions are not met, the time step is updated, and the simulation returns to the first action as described in Table 2. As the current version of the model is not intended to assess long-term population dynamics nor the establishment of an enzootic YFV circulation, the maximum time allowed for the simulation is 1825 days (5 years).

#### ODD- Additional Information

**Text A1**. The presumed maximum daily flight distance for mosquitoes *Haemagogus*.

Service (1997) pointed out that short-distance dispersion is undoubtedly the norm in mosquito biology. It is assumed that mosquitoes that disperse long distances will only do so if carried by the wind or in the absence of resources such as hosts, breeding sites, mates, resting places, and nectar sources. Little dispersion is needed if these resources can be found close to the individual. Verdonschot and Besse-Lototskaya (2014) reviewed 460 articles providing quantitative information about the flight distance of mosquitoes. Based on mark-recapture experiments, they presented data from 81 works covering 35 species (8 genera) regarding the average flight distance (in meters). They concluded that the average flight ranges varied between 25m and 6 km and depended on the species. The only work to evaluate the flight capacity and dispersal of *Haemagogus* mosquitoes was carried out by Causey et al. (1950). Table A1 presents data from three mark-recapture experiments performed by the authors in forest fragments of Passos, Minas Gerais, Brazil.

**Table A1**. Flight distance of two *Haemagogus* species obtained through three mark-recapture experiments performed by Causey et al. (1950) in the municipality of Passos, Minas Gerais, Brazil. In parentheses the old names given to the species, as they appear in the article.

| Species | Experiment | Maximum distance (km) | days | km/days |
|---|---|---|---|---|
| *H. leucocelaenus* (*Aedes leucocelaenus*) | 1 | 2 | 5 | 0.40 |
| | 2 | 5.7 | 3 | 1.90 |
| | 3 | 5 | 11 | 0.45 |
| *H. janthinomys* (*Haemagogus spegazzinii*) | 1 | 1.8 | 3 | 0.60 |
| | 2 | 4.1 | 2 | 2.05 |
| | 3 | 11.5 | 10 | 1.15 |

As observed in table A1, the presumed maximum distance traveled by *H. leucocelaenus* in a day was 1.9km, and for *H. janthinomys* was 2.05km. That information was considered to assign the default value of 20 patches (2km) for the parameter `maximum-distance-link` assuming a variation of 30% (1.4 – 2.6km) in the landscape-dependent scenario.

**Text A2**. Calibrating parameters *lv1* and *lv2*.

As explained in the section 7.2 of the manuscript, the higher the values of `lv1` and `lv2`, the greater the number of individuals that will survive and become adults in the simulation. With more adults emerging, the carrying capacity of the breeding site is reached more often, causing more immature individuals to remain in the breeding site for longer and delaying their development into adults. That is a characteristic forced by the model rules and this means that `lv1` and `lv2` affect the average minimum and maximum duration of larval development. Therefore, these two parameters were adjusted to produce realistic patterns based on empirical data on the time it takes for development from immatures to adult form, such that the development time is shorter during favorable seasons and longer during unfavorable seasons. Three values were tested for each parameter (`lv1` = 1, 1.5 or 2 and `lv2` = 5, 7.5 or 10) along with their combinations. For each combination we simulated a period of a year to calculate the montlhy average minimum and maximum development time of immatures from data obtained of each node. The results are presented in table A2.

**Table A2**. Simulated average minimum (Min-aver) and maximum (Max-aver) development time in days for immatures, based on different combinations of values for the parameters `lv1` and `lv2`.

| Parameter values | Min-aver | Min-aver-sd | Max-aver | Max-aver-sd |
|---|---|---|---|---|
| lv1=1_lv2=5 | 12.0 | 0.2 | 34.8 | 3.4 |
| lv1=1_lv2=7.5 | 12.4 | 1.0 | 43.8 | 3.6 |
| lv1=1_lv2=10 | 14.9 | 1.7 | 51.1 | 3.8 |
| lv1=1.5_lv2=5 | 12.0 | 0.2 | 37.1 | 3.8 |
| lv1=1.5_lv2=7.5 | 12.6 | 1.2 | 46.0 | 4.0 |
| lv1=1.5_lv2=10 | 15.3 | 1.8 | 53.1 | 4.2 |
| lv1=2_lv2=5 | 12.0 | 0.3 | 39.6 | 4.2 |
| lv1=2_lv2=7.5 | 12.7 | 1.4 | 48.1 | 4.3 |
| lv1=2_lv2=10 | 15.6 | 1.9 | 55.1 | 4.4 |

Table A2 reveals that while there is a minor variation in the outputs for minimum development time, there is a more pronounced variability for the maximum values, which tend to increase as the combined parameter values increase. We ultimately decided on the `lv1` = 1 and `lv2` = 7.5 combination since the minimum average corroborates with the findings of Alencar et al. (2008), and the maximum average is still feasible given that certain mosquito species, such as *Haemagogus* and other Aedini, can remain in the egg stage for extended periods during colder, less rainy periods due to their resistance to desiccation and ability to enter diapause (Marcondes and Alencar, 2010). Figure A1 provides insight into the seasonal variation in development time and its correlation with the abundance of adult mosquitoes in the simulated landscape dynamics.

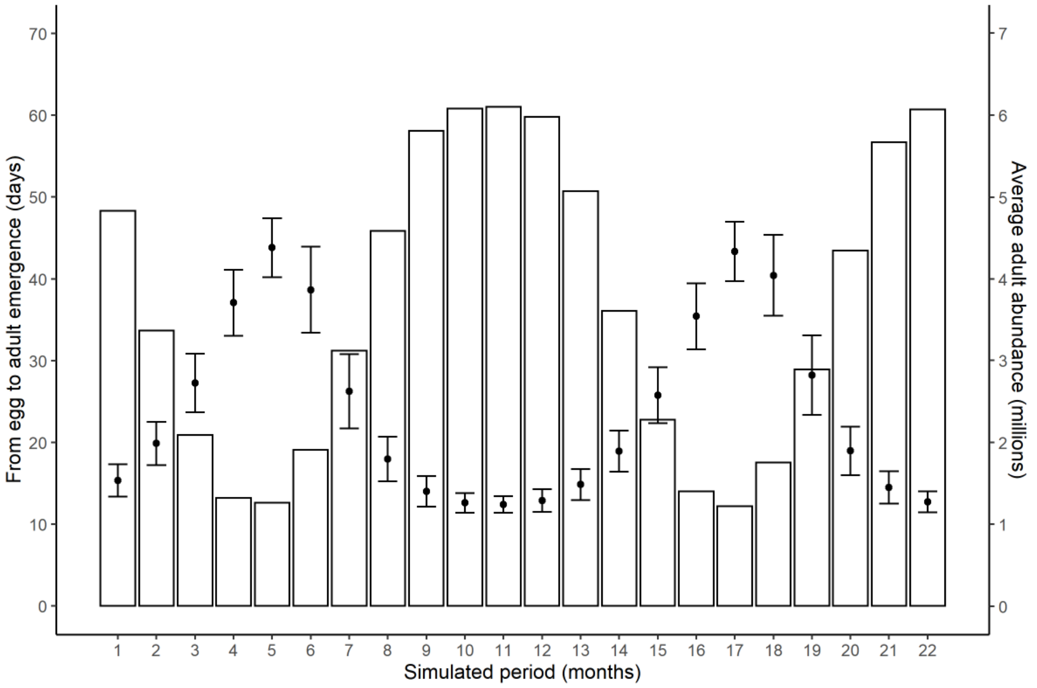


**Figure A1**. Simulated seasonal variation for average (points) and standard deviation (error bars) time from egg to adult development for mosquitoes. The bars represent the average monthly abundance of mosquitoes in the landscape. Parameters `lv1` and `lv2` were set to 1 and 7.5, respectively.

**Text A3**. Mosquitoes and hosts transmission competence for YFV.

An experimental study carried out by Couto-Lima et al. (2017) evaluated the susceptibility of *Hg. leucocelaenus* and *Sabethes albiprivus* to two South American (74018-1D and 4408-1E) and one African (S79) strains of YFV. Mosquitoes fed on infected blood at a title of 106 PFU/mL and one of the parameters measured in the study was the transmission efficiency (TE), which is the proportion of mosquitoes with virus in the salivary glands among the initial number of females tested. Data were evaluated for a period of 7 to 21 days post-feeding. The species *Hg. leucocelaenus* showed TE between 10.8 and 20% during the experiments, while *Sa. albiprivus* showed TE between 23.3 and 31.7%. Based on these values, a transmission competence of 20% was assumed as reasonable for the vector in the model simulations ($\delta_m$= 0.2).

To parameterize the transmission competence of the main host and the alternative host, data obtained by Mares-Guia et al. (2020) and Fernandes et al. (2021), who evaluated the susceptibility of new-world nonhuman primates to YFV, were considered. Both studies performed RT-PCR to identify viral load in infected individuals of different species. It was observed that for specimens of howler monkeys (*Alouatta* spp.) the distribution of Cq-values had a higher density between 8 and 15, whereas for marmosets (*Callithrix* spp.) these values tended to be above 30. Considering a standard curve with YFV vaccine 17D, a titer of $10^5$ PFU/mL had a mean Cq = 16, and 1 PFU/mL had a mean Cq = 35.6 (Fenandes et al., 2021). Thus, the value of PFU $10^6$, used in the experiments by Couto-lima et al. (2017), was considered to represent a good approximation of the viral load normally found in infected howler monkeys. Therefore, the proportion of mosquitoes with infected body among the engorged mosquitoes (IR) was used as a measure to parameterize the transmission competence of main hosts in the model simulations. Couto-lima et al. (2017) obtained IR values ranging from 48 to 70% for *Hg. leucocelaenus* and based on these values, a competence of 60% ($\delta_P$= 0.6) was assumed for main hosts. In turn, the value considered for alternative hosts was 20% ($\delta_P$= 0.2), assuming that they have a much lower viral load than the main host.

**Text A4**. The probability of finding a host for blood seeking mosquitoes.

In order to ensure that the daily probability of a blood seeking mosquito finding a host increases as more hosts are present in the area, as well as the abundance of mosquitoes goes to 0 in the absence of hosts, the model was parameterized to consider the daily probability of finding a host ($p_h$) as a function of the number of hosts in the environment.

To parameterize this probability a metamodeling strategy using the yellow fever ABM proposed by Medeiros-Sousa et al (2022) were performed, The average time taken by blood-seeking mosquitoes to find and approximate to a host before the interaction (bite) was used as output. The variation in $p_h$was obtained by vary the number of hosts in the environment from 0 to 1 individual per hectare (intervals of 0.1) and abundance from 2 to 22 mosquitoes per hectare (intervals of 2) was used to test a possible covariate effect of mosquito density. Thus 121 different combinations of host and mosquito abundance were simulated. Other parameters were set to default following the values presented in Table 3.

The simulations were performed for 3,400 steps, excluding the first 400 steps for allow model dynamics stabilization.

After obtaining the outputs, we first built linear models to explore a possible interactive effect of the two predictor variables (host and mosquito abundance) on the outputs values. Both the interactive effect ($p$ = 0.78) and the addictive effect of the variable mosquito abundance ($p$ = 0.85) were not significant, and only the variable host abundance was retained in the model ($p$ < 0.001). As the outputs tended to present an assymptotic behavior, we opted to modeling the response of $p_h$to host density using a Michaelis-Menten fit, which showed superior fit to the data when compared to a simple linear model (Figure A2). Thus the two model parameters (d: 2.38232 and e: 107.12518) can be used to calculate $p_h$ as follow: $p_h = 2.382 * \frac{Total\ hosts}{107.125 + Total\ hosts}$.

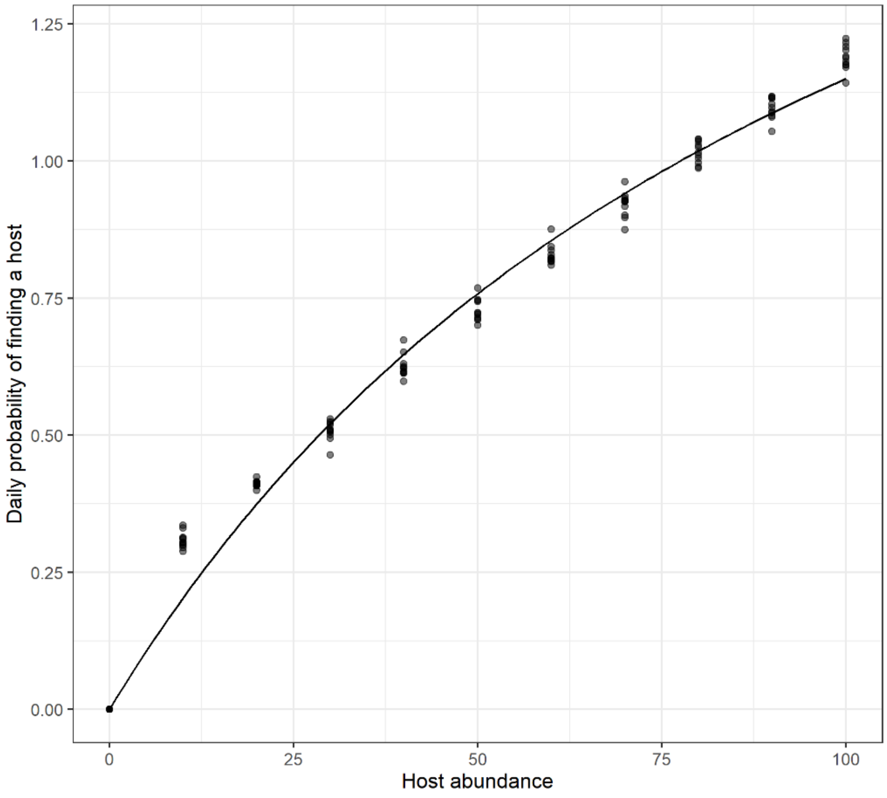


**Figure A2**. Relationship between mosquito daily probability of finding a host and the host abundance. The line represents a Michaelis-Menten metamodeling fit for output data obtained via simulations (black points).

### References (ODD- Additional Information)


Alencar, J., de Almeida, H. M., Marcondes, C. B., & Guimarães, A. É. (2008). Effect of multiple immersions on eggs and development of immature forms of *Haemagogus janthinomys* from South-Eastern Brazil (Diptera: Culicidae). **Entomological News**, 119(3), 239-244.

Causey, O. R., Kumm, H. W., & Laemmert Jr, H. W. (1950). Dispersion of Forest Mosquitoes in Brazil: further studies. **American Journal of Tropical Medicine**, 30(2), 301-12.

Couto-Lima, D., Madec, Y., Bersot, M. I., Campos, S. S., de Albuquerque Motta, M., Dos Santos, F. B., ... & Failloux, A. B. (2017). Potential risk of reemergence of urban transmission of Yellow Fever virus in Brazil facilitated by competent *Aedes* populations. **Scientific reports**, 7(1), 1-12.

Fernandes, N. C. A., Guerra, J. M., Díaz-Delgado, J., Cunha, M. S., Iglezias, S. D., Ressio, R. A., ... & Catão-Dias, J. L. (2021). Differential yellow fever susceptibility in New World nonhuman primates, comparison with humans, and implications for surveillance. **Emerging infectious diseases**, 27(1), 47.

Marcondes, C. B., Alencar, J. (2010). Revisão de mosquitos *Haemagogus* Williston (Diptera: Culicidae) do Brasil. **Revista Biomedica**, 21, 221-238

Mares-Guia, M. A. M. M., Horta, M. A., Romano, A., Rodrigues, C. D., Mendonça, M. C., Dos Santos, C. C., ... & de Filippis, A. M. B. (2020). Yellow fever epizootics in non-human primates, Southeast and Northeast Brazil (2017 and 2018). **Parasites & Vectors**, 13(1), 1-8.

Medeiros-Sousa, A. R., Laporta, G. Z., Mucci, L. F., & Marrelli, M. T. (2022). Epizootic dynamics of yellow fever in forest fragments: An agent-based model to explore the influence of vector and host parameters. **Ecological Modelling**, 466, 109884.

Service, M. W. (1997). Mosquito (Diptera: Culicidae) dispersal—the long and short of it. **Journal of medical entomology**, 34(6), 579-588.

Verdonschot, P. F., & Besse-Lototskaya, A. A. (2014). Flight distance of mosquitoes (Culicidae): a metadata analysis to support the management of barrier zones around rewetted and newly constructed wetlands. **Limnologica**, 45, 69-79.
